# The hidden elasticity of avian and mammalian genomes

**DOI:** 10.1101/081307

**Authors:** Aurélie Kapusta, Alexander Suh, Cédric Feschotte

## Abstract

Genome size in mammals and birds shows remarkably little interspecific variation compared to other taxa. Yet, genome sequencing has revealed that many mammal and bird lineages have experienced differential rates of transposable element (TE) accumulation, which would be predicted to cause substantial variation in genome size between species. Thus, we hypothesize that there has been co-variation between the amount of DNA gained by transposition and lost by deletion during mammal and avian evolution, resulting in genome size homeostasis. To test this model, we develop a computational pipeline to quantify the amount of DNA gained by TE expansion and lost by deletion over the last 100 million years (My) in the lineages of 10 species of eutherian mammals and 24 species of birds. The results reveal extensive variation in the amount of DNA gained via lineage-specific transposition, but that DNA loss counteracted this expansion to various extent across lineages. Our analysis of the rate and size spectrum of deletion events implies that DNA removal in both mammals and birds has proceeded mostly through large segmental deletions (>10 kb). These findings support a unified ‘accordion’ model of genome size evolution in eukaryotes whereby DNA loss counteracting TE expansion is a major determinant of genome size. Furthermore, we propose that extensive DNA loss, and not necessarily a dearth of TE activity, has been the primary force maintaining the greater genomic compaction of flying birds and bats relative to their flightless relatives.

## INTRODUCTION

The nature and relative importance of the molecular mechanisms and evolutionary forces underlying genome size variation has been the subject of intense research and debate [1–5]. Variation in genome sizes may not always occur at a level where natural selection is strong enough to prevent genetic drift to determine their fate (neutral or effectively neutral variation) [3]. On the other hand, a number of correlative associations between genome size and phenotypic traits, such as cell size [6, 7] and metabolic rate associated with powered flight [8, 9], suggest that natural selection and adaptive processes also shape genome size evolution. Teasing apart the relative importance of these two forces (drift and selection) requires a better understanding of the mode and processes by which DNA is gained and lost over long evolutionary periods in different taxa. Thus, establishing an integrated view of the contribution of gain and loss of DNA to genome size variation (or lack thereof) remains an important goal in genome biology [e.g. 1, 2, 10, 11].

Most studies of genome size evolution have focused on taxa with extensive variation in genome sizes, such as flowering plants [12–17], conifers [18], insects [19–22], teleost fishes [23], or species with extreme sizes (such as pufferfishes [24, 25] and salamanders [26, 27]). Together, these investigations have documented that the differential expansion, accumulation, and removal of transposable element (TE) sequences represent a major determinant of genome size variation in plants and animals [see for reviews 2, 5, 28]. Generally, the studies cited above have revealed that species with larger genomes tend to have larger TE content combined with low rates of TE DNA removal, and *vice versa* for smaller genomes.

In comparison to these taxa, birds and mammals show little interspecific variation in genome size (from ~1 to 2.1 Gb and 1.6 to 6.3 Gb respectively, Figure 1), and little is known about the mechanisms underlying genome size homeostasis in these two classes of amniotes. In contrast to plants (and to a lesser extent fishes), changes in ploidy do not appear to represent a major source of genome size variation in birds or mammals, and there is no evidence of whole genome duplication events during amniote evolution [29]. On the other hand, it is well established that a considerable amount of new nuclear DNA has been generated throughout eutherian and avian evolution, mostly via TE expansion and, to a lesser extent, through segmental duplications [30–41]. These observations thus raise a conundrum whereby TE activity has been pervasive in mammalian and avian evolution, yet has had apparently little impact on genome size.

**Figure 1:**
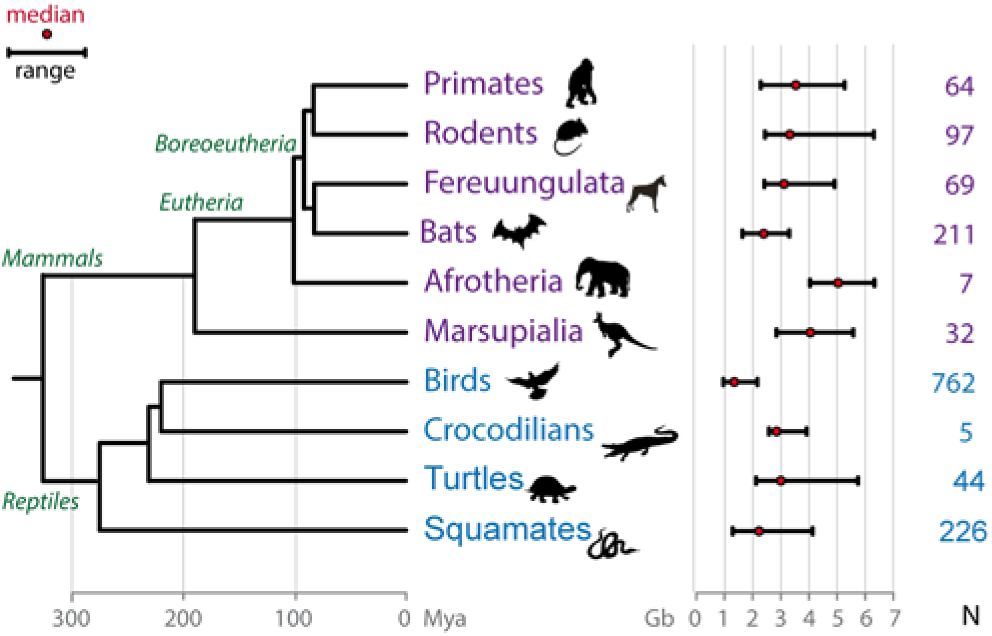
Genome size variation in amniotes. Genome size ranges of different groups of species are shown as black bars (from smallest to largest genome sizes). Birds range from ~1 to 2.1 Gb and mammals from 1.6 to 6.3 Gb, which is in contrast to the range among all vertebrates (~0.4 to ~133 Gb) [114]. Divergence times are represented on the phylogenetic tree in My as in [115, 116]. Red dot: median. N: numbers of species inside each group with genome size data included in the figure (compiled from [114] as of March 6^th^ 2015). When several measures exist for one single species, values are averaged. Gb: gigabases. For rodents, the red viscacha rat was not included (tetraploid, genome size estimated to 8.4 Gb).

The simplest way to reconcile this conundrum is to postulate that the amount of lineage-specific DNA gained by transposition has been systematically equalized by the removal of DNA along those lineages, thereby accounting for genome size homeostasis in mammals and birds [as hypothesized in 11, 42]. However, this hypothesis remains largely untested and, overall, little is known about the mode and tempo of genomic DNA loss in amniotes. An earlier comparative genomic study of insertion/deletion (indel) rates across 13 vertebrate genomes [43] implicated that variation in DNA gains through TE expansion acted in concert with variations in deletion rates to modulate genome size during evolution, but the region analyzed was limited to a 12-Mb alignment and included only a single bird species. A more comprehensive analysis of DNA gain and loss in the lineages of human, mouse and dog [33] showed that the dog and human lineages experienced 2.5x less DNA loss than in the mouse lineage, but also 2.8x and 1.6x less DNA gain, a balance explaining the modest differences in genome size across these species. Little is known about genome size dynamics in birds. Statistical models of genome size evolution in the avian lineage have inferred that a drastic contraction event (by ~0.8-fold) occurred prior to the divergence of birds in a theropod ancestor [44]. Consistent with this idea, a recent comparative analysis of 48 bird genomes revealed that introns and intergenic regions are, on average, ~2x and ~1.4x smaller in birds than in mammals and non-avian reptiles, respectively [40] [see also 45, 46]. Furthermore, the ostrich lineage was found to have experienced, on average, larger genomic deletions than alligator and turtle lineages [40]. Despite these recent insights, our understanding of genome size dynamics across eutherian and avian evolution remains fragmentary. In particular, the mode and tempo of DNA loss throughout amniote evolution has not been examined systematically.

Leveraging the recent sequencing of dozens of mammalian [e.g 47] and avian [40] genomes, we characterize genome size evolution in mammals and birds through an integrated analysis of DNA gain and loss on a genome-wide scale. The results are consistent with an ‘accordion’ model of genome size homeostasis whereby DNA gains are balanced by DNA loss, primarily through large-size deletions.

## RESULTS

### Genome size evolution as the integration of gain and loss of DNA

We used available genome assemblies for 10 eutherian (placental) mammals and 24 avian species, and their respective TE annotation (Methods). We first estimated the amount of DNA gained via transposition events in each lineage since their last common ancestor. For mammals, we compiled data from the literature as well as our own analyses to divide TE families previously characterized in each species into lineage-specific and ancestral families (Methods and Table S1). Using this information, we applied the RepeatMasker software [48] to infer the amount of DNA occupied (and therefore gained) by lineage-specific TEs in each of the genome assemblies examined. We added the amount of lineage-specific DNA gained by segmental duplications, when documented in the literature (information limited to some mammals) [31, 34, 35, 37, 38]. Because the evolutionary history of bird TE families has not been characterized as extensively as in mammals, we inferred the age of each TE insertion based on its divergence to the cognate family consensus sequence using lineage-specific neutral substitution rates [40] (Methods). For birds, gains were estimated from the DNA amounts corresponding to insertions younger than 70 My, which corresponds to the onset of the Neoaves radiation [49].

We then computed the total amount of DNA lost in each lineage by subtracting the amount of ancestral genomic DNA of each species (assembly size minus gains) to the “projected” assembly size of their common ancestor. It is important to note that our analysis requires using assembly sizes, which is the genomic space where TEs have been annotated, rather than actual genome sizes (which are always slightly larger due to current limitation in assembling highly complex regions such as centromeres). For eutherians, we used an ancestral genome assembly size previously estimated at 2.8 Gb based on a multiple alignment of 18-species genome assemblies, allowing ancestral reconstruction [50, 51]. For birds, we used 1.3 Gb as the predicted assembly size for both the ancestor of Paleognathae and Neoaves based on ancestral genome sizes previously inferred for these two clades [52] and a comparison of genome sizes and assembly sizes for each of the bird species sequenced (Methods). Using these inferences, we estimated the total amount of DNA lost along each of the 34 lineages considered (Dataset S1).

As an example of the approach, in the human lineage we estimated that 899 Mb of the hg38 assembly consisted of DNA gained via lineage-specific TE insertions (815 Mb) and segmental duplications (84 Mb), which leaves an “ancestral DNA” amount in the human genome assembly of 2,150 Mb. Thus, we can infer that 650 Mb (2,800 minus 2,150) of ancestral DNA was lost in the human lineage over the past ~100 My. The same procedure was applied to the other species lineages considered. Based on these amounts, we computed lineage-specific DNA loss coefficients *k* [as in 33] with *E = A e*^*-kt*^, where *E* is the amount of extant ancestral sequence in the species considered, *A* the total ancestral assembly size, and *t* time (100 My for mammals and 70 My for birds) (Methods and Dataset S1). Applying our predicted DNA loss rate for human, we obtain a coefficient *k* of 0.0026 per My, which is nearly identical to the previously calculated coefficient of 0.0024 [33], despite being based on a different methodology to infer DNA gain and loss.

When applied to all lineages (Figures 2, 3 and S1), the results of these analyses show that the amount of DNA gained and lost has varied substantially across lineages. DNA gains vary by more than 6-fold across mammals (from 150 Mb in the megabat to 1,007 Mb in the mouse lineages, Figure 2) and by more than 30-fold across birds (from 7 Mb in the ostrich to 255 Mb in the woodpecker lineages, Figure 3). DNA loss amounts range by 2-fold across mammals (from 650 Mb in the human to 1,373 Mb in the microbat lineages), and by more than 3-fold across birds (from 119 Mb in the ostrich to 424 Mb in the woodpecker lineages).

**Figure 2:**
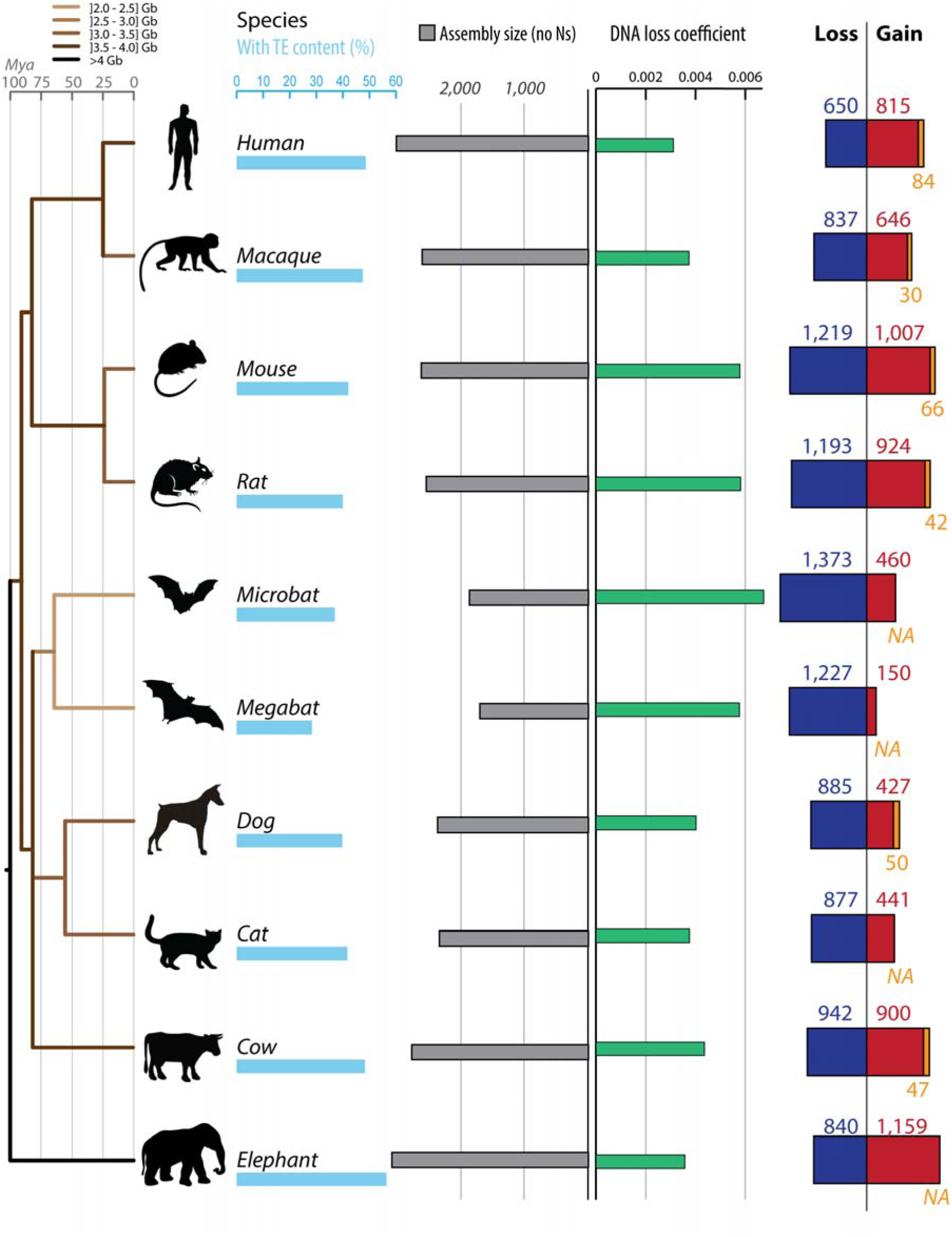
Gain and loss of DNA in 10 mammalian lineages. For each species, phylogenetic relationship (left panel) [116], TE content (light blue bars), assembly sizes (with N removed, grey bars), DNA loss coefficients (green bars) as well as gain (red and orange) and loss (dark blue) of DNA are shown. DNA gains correspond mostly to lineage-specific TEs in red (not shared with other mammals). When available, measures of segmental duplications (SDs) were added (orange). Because SDs also contain TEs, we corrected the SD amounts with the TE content of each genomes. DNA loss amounts and rates are calculated as in [33] using a common ancestor ‘assembly’ size for all mammals of 2,800 Mb (Methods). Phylogenetic tree is color-coded based on genome sizes in pg (combination of data from [114]) based on parsimony. See Dataset S1 for numbers, calculation steps and assemblies details. All numbers are in Mb.

**Figure 3:**
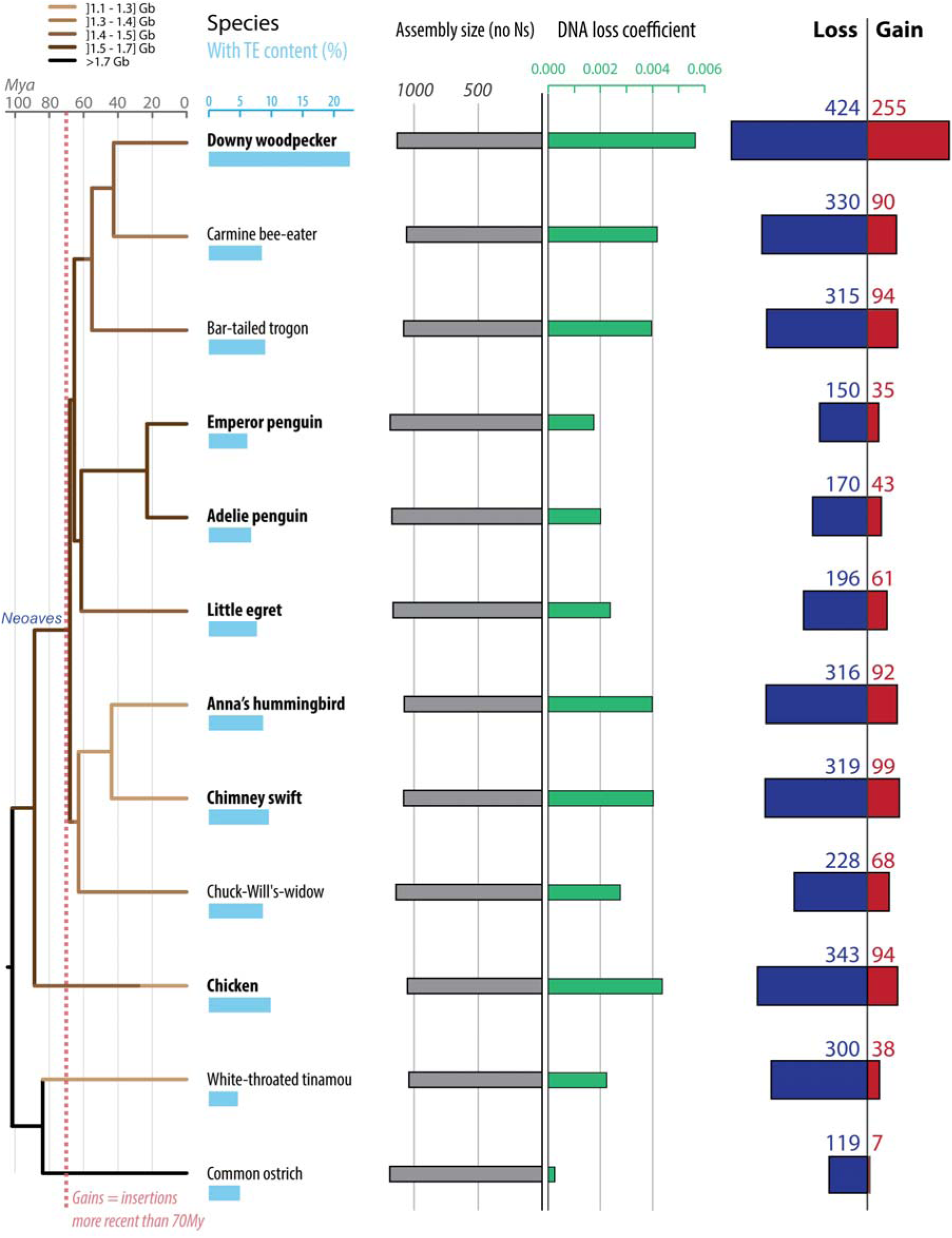
Gain and loss of DNA in 12 avian lineages. For each species, phylogenetic relationship (left panel) [49], TE content (light blue bars), assembly sizes (with N removed, grey bars), DNA loss coefficients (green bars) as well as gain (red) and loss (dark blue) of DNA are shown. Species names in bold correspond to high coverage genomes, and the others to low coverage genomes [40]. DNA gain corresponds to insertions younger than 70 My. DNA loss amounts and rates are calculated as in [33] using a common ancestor size of 1,300 Mb (see text and Methods). Phylogenetic tree is color coded based on genome sizes in pg (combination of data from [114] and extrapolations from assembly sizes and coverage), based on parsimony as in [52]. See Dataset S1 for numbers, calculation steps and assemblies details. All numbers are in Mb.

For mammals, these results confirm the trends previously reported for some of these lineages [30, 31, 33]: we observe more gains in rodents than in human, and more gain in human than in dog, together with more loss in rodents than in dog or human. In addition, we found that the coefficient at which DNA was lost along the lineages examined is the lowest in the ostrich lineage and the highest in the microbat lineage (Figures 2 and 3). In fact, both the microbat and megabat lineages stand out as having both the highest amount and coefficient of DNA loss, followed closely by the mouse and rat lineages (Figures 2 and 3). Altogether, we found that neither DNA gain nor loss can solely explain variation in genome (assembly) sizes among the mammals and birds examined (Dataset S1). These results imply that variations in DNA gain and loss have acted in concert to modulate genome size in eutherian and avian evolution. We note that loss exceeded gain in all but two lineages (human and elephant, Figures 2, 3 and S1).

To investigate the extent by which these two opposite forces each contribute to genome size homeostasis, we next examine whether DNA gains (percentage of the ancestor assembly size) and DNA loss coefficients correlate with assembly sizes. In mammals, we observe a significant positive correlation between DNA gains and assembly sizes, while DNA loss are negatively (but non-significantly) correlated with assembly sizes (R^2^ = 0.73 with *p* = 0.02 and R^2^ = 0.49 with *p* = 0.15, Dataset S1). However, these results have to be interpreted with caution because our statistical power is limited by the relatively small sample of mammal lineages examined (n=10). For the 24 bird lineages analyzed, we observe that DNA loss coefficients, but not DNA gains, significantly correlate with assembly sizes (R^2^ = 0.63 with *p* = 0.0005 and R^2^ < 0.1 respectively). This observation indicates that DNA loss is a major contributor to genome size homeostasis in birds. However, neither one of these two forces alone can fully account for the variation between extent assembly sizes (e.g woodpecker, Dataset S1). Thus, these two forces must have acted in concert to modulate genome size throughout avian evolution. Consistent with this idea, we found that DNA gains and DNA loss coefficients are positively and significantly correlated with each other across the bird lineages examined (Pearson coefficient R^2^ = 0.65, *p* = 0.0005, Figure 4). These data support a model where genome size homeostasis is maintained through DNA loss counteracting DNA gains through TE expansion. This is most strikingly illustrated in the woodpecker lineage, which shows both the largest amounts of gains and the highest DNA loss coefficients (Figures 3 and S1, Dataset S1).

**Figure 4:**
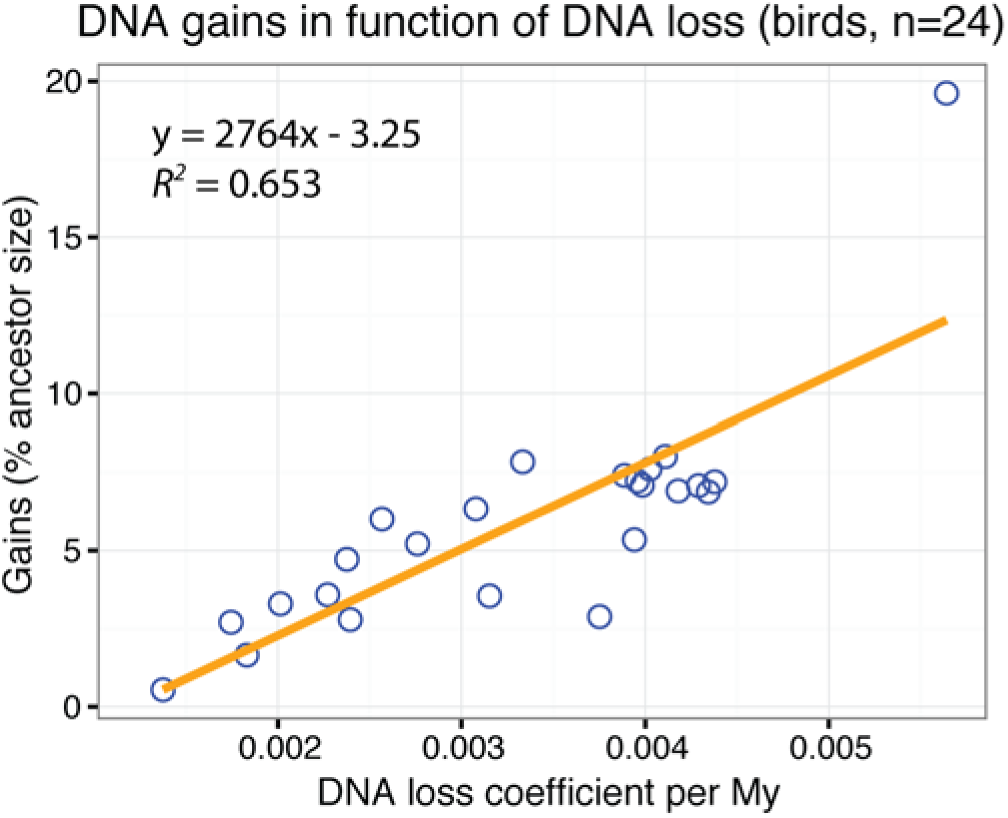
Gain and loss of DNA as driving forces of genome size variation. DNA gains are plotted in function of DNA loss coefficients for the 24 birds examined. With R^2^ = 0.658 (r^2^ on graph) and n = 24, *p* = 0.000474. The values and R [117] command lines used to build this figure can be found in Dataset S1.

### Contribution of microdeletions to DNA loss

Having determined that DNA loss makes an important contribution to eutherian and avian genome size homeostasis, we next sought to investigate the types of deletion events involved in the process. We first assessed the impact of small deletions (< 30 bp; hereafter ‘microdeletions’) through multispecies alignment available from the UCSC genome browser (MultiZ output of 100-species alignments). We separately extracted and analyzed genomic alignments for 11 eutherian species (plus the marsupial *Monodelphis domestica* as outgroup) and for seven bird species (plus the lizard *Anolis carolinensis* as outgroup). We used the principle of parsimony to infer and place microdeletion events (estimated as sequence gaps of less than 30 bp) on the phylogenetic tree of the species (Methods). The total amount of concatenated alignment analyzed in this way corresponded to 237 Mb and 52 Mb for mammals and birds, respectively. Microdeletion rates were obtained for each lineage by normalizing the amount of gaps in the alignment by its total length and dividing the amount of alignment gaps per lineage by the corresponding branch length in My (Figure 5).

**Figure 5:**
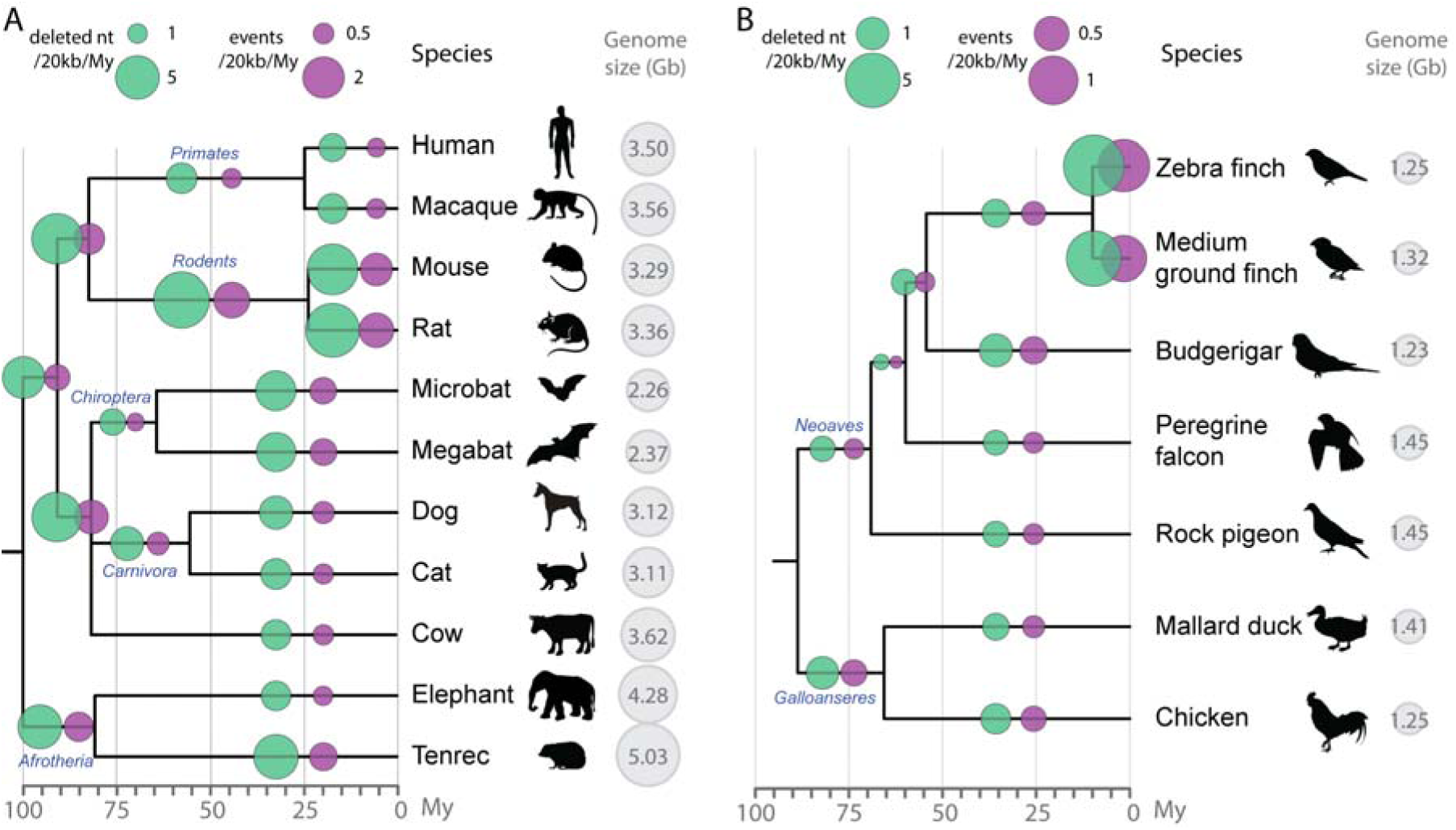
Microdeletion rates across amniotes. Microdeletion (1-30 bp) rates and number of events are shown in green and purple respectively. Microdeletions are estimated from gaps in the UCSC MultiZ 100-species alignment of human chromosomes 1 to 22, restricted to blocks containing information for the species studied (Methods and Dataset S2). Deletion rates are calculated by dividing the amount of gaps specific to each branch (not present in any other species) by My of each branch. My: million years. Gb: gigabases. Genome sizes are from [114]. There were no reported genome sizes for three species, so the size of the closest species is shown: the Tenrecidae *Setifer setosus* for *E. telfari*, *M. fuscata* for *M. mulatta*, and the average of two birds of the same family (Emberizidae) for the medium ground finch. Scales are indicated on top (note the difference between panels A and B). A. Microdeletion rates in 11 placental mammals (with *Monodelphis domestica* as outgroup). The total length of the alignment is 297 Mb and timescales are as in [31, 116, 118, 119]. Names of orders are indicated on the tree (in blue). B. Microdeletion rates in seven birds (with *Anolis carolinensis* as outgroup). The total length of the alignment is 66.5 Mb. Timescales are from [49]. Names of two super-orders are placed on the tree (in blue).

The results show that rodents have the highest microdeletion rates among the lineages examined (Figure 5A), about 3.5x higher as those for the human lineage, in agreement with previous analyses of a smaller number of mammals [26, 30, 31, 53, 54]. All other mammal lineages we analyzed exhibit microdeletion rates that are intermediate between those of human and rodents, except for the common ancestor of bats (Chiroptera) which displays the lowest microdeletion rate in our analysis. Overall, microdeletion rates do not appear to vary substantially within a given mammalian (super)order (Primates, Rodentia, Chiroptera, Carnivora). One exception is Afrotheria, where the elephant lineage is characterized by a much lower (0.4x) microdeletion rate than that of the tenrec (*E. telfari*) or that of their common ancestor. This could be linked with peculiar characteristics evolved in the elephant lineage, such as large body size, long gestation, slow development to maturity, long generation time [see for review 55]. When the number of microdeletion events (normalized for alignment length) is considered, rather than the total amount of DNA removed by microdeletions (Figure 5A), we observe similar trends whereby bat, primate, and rodent lineages display the lowest, second lowest, and highest microdeletion rates, respectively.

Among the seven bird species considered in the alignment, we found that the two finches display the highest microdeletion rates (both in amount of DNA removed and number of events), which are 5.4x higher as those in the falcon lineage. All other bird lineages show rates and number of events that are intermediate between those of finches and falcon. We find that the average length of microdeletions in birds (5.8 bp) is slightly larger than inferred in mammals (5.2 bp) (Dataset S2). Average lengths per species lineages range from 4.8 bp (bats) to 5.7 bp (tenrec) in mammals and from 5.5 bp (zebra finch) to 6.1 bp (rock pigeon) in birds, and the distribution profiles are significantly different in pairwise species comparisons between birds and mammals, except between the zebra finch and some of the mammals (Dataset S4 and Figure S2). Within orders, pairwise species comparisons are significantly different between bats and other mammals, and between zebra finch and the other birds (Dataset S4 and S2). Microdeletions are slightly smaller in size in human and macaque than in mouse and rat (by 0.46 bp on average), which is in agreement with a previous study [54]. In summary, we observe substantial variation in the rate and size spectrum of DNA microdeletion between and within eutherian and avian orders. This may reflect species characteristics such as mechanisms at the origin of microdeletions and effective population size [56] (see Discussion).

Next, we applied the microdeletion rates inferred over the various branches of the phylogeny to extrapolate the amount of DNA lost through this class of deletion during the last 100 My in mammals and the last 70 My in birds. We compared this to the total amount of DNA estimated to be lost within the same timeframe (Figures 2, 3 and S1). The results indicate that microdeletions account for only a small fraction (<10%) of the DNA lost in mammals and birds over the past 100 My and 70 My respectively (from 1% for the chicken lineage to 8.2% in the rat lineage; Dataset S2).

### Contribution of midsize deletions to DNA loss

The method above enables the extrapolation of microdeletion rates over the various branches of the tree, but relies on a multispecies alignment using human as a reference. Therefore, it inherently favors the retention of regions alignable between human and the species considered, potentially biasing our analysis for the most conserved regions of the genome. Additionally, large deletions are, by design, excluded from the MultiZ alignment blocks used in the analysis above. In order to estimate microdeletion rates in a less biased fashion, as well as to capture larger deletion events, we developed an independent approach relying on the comparison of closely related species. We designed our computational pipeline to capture deletions up to a specified length (10 kb) in trios of species representative of the primate, chiropteran (bats), carnivore, artyodactyl, and afrotherian lineages, as well as eight trios of bird species (Figure 6). Briefly, the approach selects at random a pair of anchor sequences separated by a set length in the genome of the outgroup species as a query in BLAT searches to identify orthologous regions in the other two species (Methods). Three-way species nucleotide sequences alignments are then generated and deletions are quantified from the amount of gaps in the alignment and parsimoniously placed along the branches of the phylogeny (Methods). Deletion rates were obtained by dividing the total length of alignment gaps per given species lineage by the corresponding branch length in My. To be able to compare across lineages, the deletion rates were also normalized by alignment length (shown in Mb in Figure 6).

**Figure 6:**
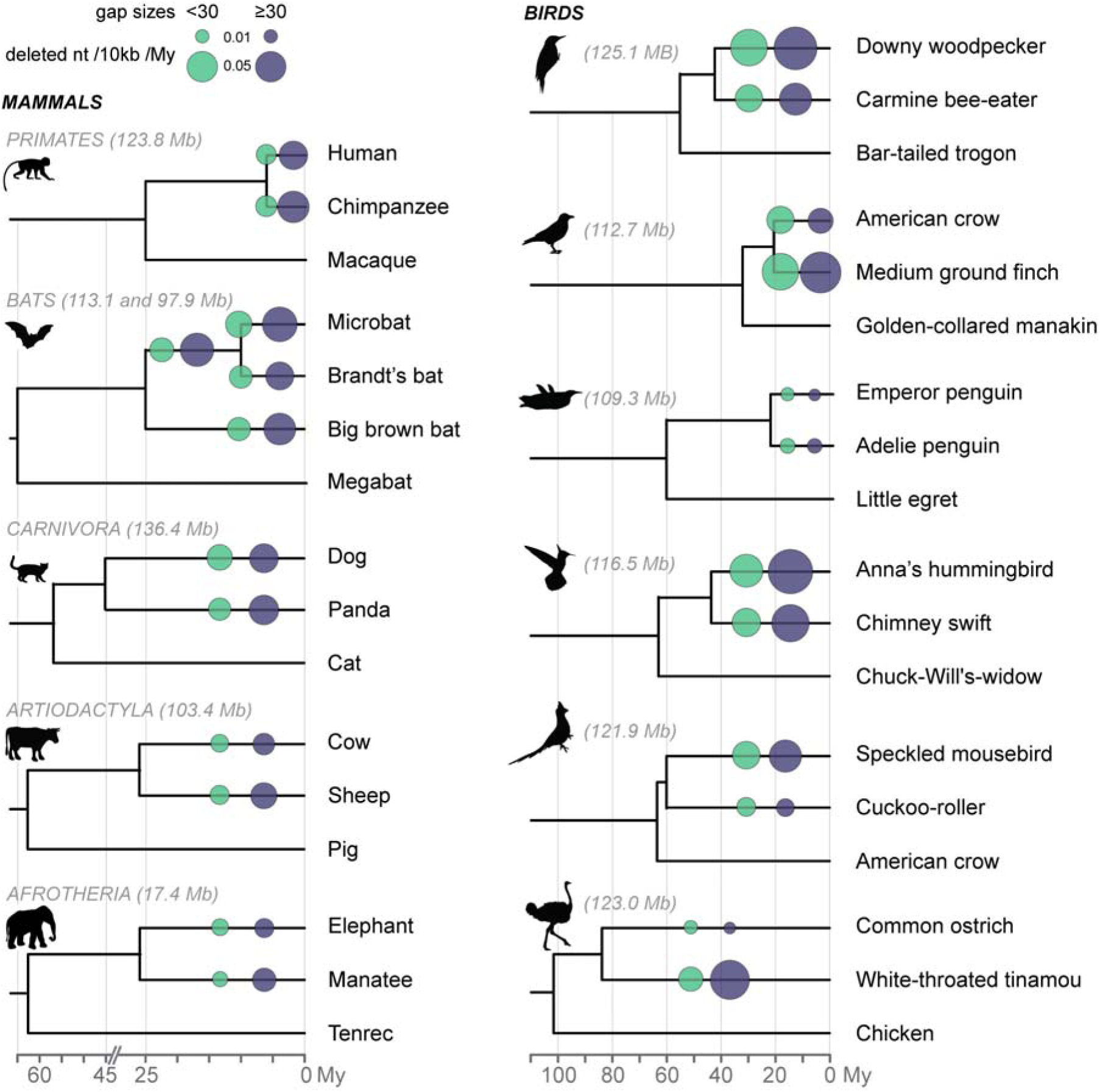
Midsize deletion rates across amniotes. Microdeletions (1 to 30 bp) and midsize deletion (from 30 bp to 10 kb for mammals and birds, respectively) rates, measured after a recent split between two species, are shown in light green and dark blue, respectively. Deletion rates were calculated based on gaps in alignments of orthologous specific to the species (Methods) and are normalized by alignment length (in Mb on the figure). We placed the deletion events on the phylogenetic tree based on parsimonious polarization via the respective outgroup species. Gaps were considered shared by multiple species when they overlapped for at least 85% of their respective lengths (Methods). For bats, 2 sets of 3 species were considered: group I with *M. lucifugus*, *M. brandtii*, and *E. fuscus* (113.1 Mb of alignment), group II with *M. lucifugus*, *E. fuscus*, and *P. vampyrus* (97.9 Mb of alignment). Rates placed on the *Myotis* branch (after the split with *E. fuscus* but before the split between *M. lucifugus* and *M. brandtii*) were inferred from *M. lucifugus* rates. See Dataset S3 for the numbers, and Figure S3 for the midsize deletion spectrums.

We first used these datasets as an independent method to infer microdeletion (<30 bp) rates along the lineages represented and compared the results with those obtained with the MultiZ approach outlined above. We observe that the trends observed using the MultiZ approach for this subset of species are largely recapitulated (Figure 6, light green circles). For instance, the elephant displays the lowest microdeletion rates among the species compared, while bats and medium ground finch display the highest. However, microdeletion rates inferred by this method are consistently higher than those estimated based on the MultiZ alignment, on average 1.6 times higher for mammals and 4.3x higher for birds (Dataset S3). Presumably this difference reflects the greater evolutionary constraint of the genomic regions aligned by MultiZ, which generally leads to an underestimation of the “neutral” microdeletion rates of the species. Nevertheless, the fraction of total DNA loss accounted for by microdeletions when applying the new rate estimates remain modest, ranging from 5.1% in the cow lineage to 15.4% in the medium ground finch lineage (Dataset S2). Thus, the vast majority of DNA loss during eutherian and avian evolution must have occurred through deletions larger than 30 bp.

We then sought to capture deletions larger than 30 bp, and an analysis of the size spectrum of alignment gaps recovered shows that our computational pipeline succeeded in capturing relatively large deletion events (Figure S3). For example, in the human lineage, 5.8% of the gaps were longer than 1 kb and the largest deletion event identified was 9,022 bp relative to the macaque genome (breakpoint in hg38 at chr10:81816824). Overall, between 10 and 23% of the deletion events recovered in each species were larger than 200 bp (Figure S3). We manually verified the longest events recovered in each species: in mammals the largest was a ~10-kb deletion in the chimpanzee relative to the macaque genome (breakpoint in panTro4 at chr14:92050068). In birds, the largest event corresponded to a ~6-kb deletion in the medium ground finch relative to the golden-collared manakin (see legend of Figure S3 for the breakpoint coordinates of all longest events). Because longer deletion events tend to be fragmented into several gaps when the sequences of the three species are aligned, counting the number of gaps in the alignment would likely overestimate the actual number of deletion events in each species lineage. Thus we focused our analysis of midsize (>30 bp) deletion events on the *amount* of DNA lost through this class of deletion, rather than the actual number of events.

The results across the five eutherian orders examined (Figure 6) reveal trends similar to our analysis of microdeletion rates, with the elephant and the microbat showing the lowest and highest rates of midsize deletions, respectively (Figure 6, dark grey circles). By applying the rates of midsize deletion inferred for each mammalian lineage to the entire distance separating these species from their common ancestor (~100 My ago), we were able to estimate that the amounts of DNA lost via this class of deletion ranged from 62 Mb in the elephant lineage to 134 Mb in the human lineage. These extrapolated figures (Dataset S3) suggest that midsize deletions have accounted for 7.3% (elephant) to 20.7% (human) of the total amount of DNA lost during eutherian evolution (11% on average). Together, micro-and midsize deletions account for about 30.9% of DNA loss in the human lineage, and only 14.1% in the microbat lineage (13% in the elephant lineage, 18% on average, Dataset S3). These data suggest that the vast majority of eutherian DNA loss has occurred through deletion events larger than those we are able to capture here (~10 kb).

For birds, the results of our midsize deletion analysis reveal trends similar to those for microdeletions as well: the medium ground finch, Anna’s hummingbird, and woodpecker lineages show the highest rates, while the smallest rates are observed in the ostrich and penguin lineages (Figure 6 and Dataset S3). Next, we sought to assess the relative contribution of micro- and midsize deletions in birds to genome size homeostasis. We took advantage of the statistical power enabled by the analysis of 16 species to test the relationship between the rates of these two classes of deletions and the DNA loss coefficients calculated over 70 My of evolution (Dataset S3). We observe that the two variables are positively correlated, either if each class of deletion is considered separately (R^2^ = 0.79 with *p* = 0.0003 and R^2^ = 0.63 with *p* = 0.009 for micro- and midsize deletions respectively, Dataset S3) or together (R^2^ = 0.73 with *p* = 0.0015, Dataset S3). These results suggest that both classes of deletions are significant drivers of genome size homeostasis during avian evolution.

The extrapolation of the amount of DNA lost in birds through the combined action of micro-and midsize deletions vary by up to one order of magnitude across lineages, from 14 Mb (ostrich and Adelie’s penguin) to 136 Mb (woodpecker) (Dataset S3). These amounts account from 9.5% (Adelie’s penguin) to 39.2% (Anna’s hummingbird) of the total DNA loss along the bird lineages examined (21.7% on average). Thus, on average, the contribution of two classes of deletions is similar in birds and mammals, but we observe a greater variation in micro- and midsize deletion rates among birds than among mammals.

## DISCUSSION

### DNA gain and loss analysis reveals the elasticity of avian and mammalian genomes

Our study represents, to our knowledge, the most systematic analysis of the amount of genomic DNA gained and lost during eutherian and avian evolution, two taxa showing remarkably little interspecific variation in genome sizes compared to others (Figure 1). One interpretation for this apparent stasis in genome size could be that these lineages simply experienced relatively small amounts of DNA gain and loss during evolution [57–59]. Our analysis shows that this is clearly not the case: there has been extensive DNA gain and loss of DNA throughout eutherian and avian evolution. For example, the amount of DNA gained via lineage-specific transposition in the mouse lineage contributed to a net gain of DNA equivalent to 33% of the current genome content, while the equivalent of 44% of genome content was lost over the same timeframe (Figure 2 and Dataset S1). The woodpecker lineage provides another striking example. Amongst birds, this species lineage has experienced the largest amount of DNA gain (255 Mb, predominantly through CR1 LINE transposition [40]) but also the largest amount of DNA loss (424 Mb, equivalent to about a third of the genome) over the past ~70 My, resulting in a current genome size comparable to that of other modern bird species (Figures 3 and S4, Dataset S1). Thus, our data reveal a previously underappreciated level of elasticity in eutherian and avian genomes.

These findings allow us to uncover a general pattern of genome evolution along the major avian and eutherian lineages, whereby the (often large) amount of DNA gained via lineage-specific transposition is essentially balanced by the amount of DNA lost over the same timeframe. This ‘accordion’ process helps explaining the relative maintenance of genome size across the eutherian and avian phylogeny. This is particularly evident in birds (Figure 1), which display a positive correlation between DNA gain and DNA loss (Figure 4). Thus, our results indicate that the relatively small genome size of birds is not merely due to a dearth of transposition in those lineages, as previously hypothesized, [e.g. 57, 58, 59], but rather the result of a dynamic interplay between TE-mediated DNA acquisition and subsequent DNA loss [as suggested in 11, 42].

### DNA loss through large deletions as a determinant of genome size

Previous studies assessing DNA loss have mainly focused on deletions within TE sequences, which imposes a relatively small upper limit for the size of observable events (since TE copies rarely exceed 10 kb). The rate of deletions estimated through this approach have been shown to be a major predictor of genome size evolution in insects [e.g. 19, 20-22, 60], plants [e.g. 13], and a few vertebrates [11, 24–27]. However, whether the variation in the rate of small deletions can actually account for genome size variation observed between taxa has been questioned [discussed in 61, 62]. Indeed, quantifications from limited comparative datasets have suggested that microdeletions alone cannot account for the extent of genome contraction observed in some vertebrate lineages [e.g. 2, 23, 53, 54, 63]. Here, we assessed a broader size spectrum of deletion through whole-genome and local alignments of diverse birds and mammals. Our estimates of microdeletion (1-30 bp) rates show that this type of events can only explain a minute fraction of the DNA content lost during avian and eutherian evolution (Figure 5 and Dataset S3) and as such do not appear to be a major contributor to genome size evolution in these taxa.

Our results show that midsize deletions (31 bp to 10 kb) play a larger role than microdeletions in explaining the observed interspecific variation in DNA loss. Collectively, however, micro and midsize deletions detected in our analyses still account for a limited fraction (9.5 to 40% and 20% in average, Dataset S3) of total DNA loss in eutherian and avian evolution. These data suggest that the vast majority of DNA loss in amniotes has been driven by relatively large deletions (>10 kb). Such large deletions are challenging to detect systematically with currently available genome assemblies, precluding us to measure the rate of these events along the lineages considered in this study. We note, however, that instances of large chromosomal deletions have been documented previously in mammals (e.g. 1,511 and 845 kb [64] and 31 kb [65]; see also [66]), and we were able to identify individual events (Figure S5). Similarly, large segmental deletions were inferred to have had occurred in the common ancestor of birds (118 events for a total of 58 Mb and up to 2.1 Mb per event [40]).

Such large deletions, combined with the sheer amount of DNA loss in some of the mammal and bird lineages examined (up to 37.9% and 22.6% of nuclear DNA content respectively), underscores the dispensability of a large fraction of genomic DNA in these animals [66–68]. Yet it does not preclude that the process of segmental DNA loss has played an important role in driving phenotypic evolution [69–71]. In fact, there are strong hints that large deletions caused a substantial level of gene loss in birds (274 protein-coding genes) with potentially profound phenotypic consequences [40, 72, 73]. The foreseeable improvement of genome assembly via third-generation sequencing (e.g. long-read sequencing and gap filling) [see 74] will provide a way to more directly test the hypothesis that large deletion events play a prominent role in amniote genome evolution.

### Genome contraction covaries with transposable element expansion

What could be the mechanisms facilitating the co-variation between the amounts of DNA gained via TE insertions and the DNA that is lost, which is especially striking in bird evolution (Figure 4)? One simple explanation would be that TE insertions and deletions occur and fix at comparable rates in a given species lineage because they are governed by the same population genetics parameters, which also govern variation in other, largely neutral mutational processes such as nucleotide substitutions [1, 56]. Consistent with this idea, we find that microdeletion rates (and, to a lesser extent, midsize deletion rates) correlate strongly with neutral substitution rates in birds (Datasets S2 and S3). This may suggest that variation in microdeletion rates largely reflects population genetic parameters, with large effective population sizes leading to an uncoupling of neutral genetic variation from nearby deleterious alleles, i.e., a less pronounced effect of linked selection [75]. Conversely, natural selection acts less efficiently in species with smaller effective population sizes [75], which has been suggested to contribute indirectly to the reduced purging of slightly deleterious TE insertions [3, 76, contra 77]. Mammals have generally smaller effective population sizes than birds [78], which is predicted to increase the probability of fixation of nearly-neutral TE insertion and deletion events through genetic drift [75, 79]. This prediction is congruent with our observation that the overall amounts of DNA gained and lost have been more substantial in mammals than in birds (Figures 2, 3 and S1).

Mechanistically, the circumstances of frequent fixation of TE insertions would also provide a plausible fodder for large chromosomal deletions. Indeed, interspersed repeats with high level of sequence similarity such as recently expanded TE families represent a prime substrate for non-allelic homologous recombination (NAHR) events that may result in the deletion of the intervening DNA (reviewed in [80]. While the impact of TE-mediated NAHR on the process of DNA loss has been well-documented in plants [e.g. 12, 15, 16, 81-84] [see for review 28], it has not been systematically examined in vertebrates. Nonetheless, comparative studies in primates have suggested that an increased density of TEs from the same family augments the probability of inter-element NAHR deletion events to occur between TE copies. For example, the highly abundant *Alu* elements have mediated considerably more NAHR deletion events [e.g 65, 85, 86] than L1 [87] or SVA [88] elements, which occur at much lower density in primate genomes. Thus, it is tempting to speculate that the explosive amplification of one or a few TE families, such as CR1 elements in woodpeckers [40] and Ves SINEs in bats [89, 90], led to an increase opportunity for NAHR thereby facilitating the extreme degree of DNA loss that we observed in these two lineages (Figures 2, 3 and S1). The idea that genome expansion via transposition subsequently promotes genome contraction via large-scale TE-mediated deletion would provide a mechanistic underpinning for the proposed ‘accordion’ model of genome size evolution.

### Implications for the origin of flight in amniotes

Overall, our findings provide support for a general trend of strong genome contraction throughout the evolution of bats and birds (Figures 2, 3, S1 and S4), the only vertebrates capable of powered flight, which extends the previous notion that the evolution of the small genomes of bats and birds predates the emergence of flight [44, 91]. The continuous genome contraction we see along multiple bird lineages (Figures 3, S1 and S4) is consistent with previous inference that their common ancestor had a larger genome than that of extant avian species [52]. Importantly, we found no significant elevation in microdeletion rates in the respective common ancestor of bats, Palaeognathae (ratites and tinamous), Galloanserae (chickens and ducks), or Neoaves (all remaining birds) (Figure 5). Paradoxically, bats display the lowest microdeletion rate in our analysis (Figure 5A). In birds, our results are in agreement with previous estimates of rates of deletions <100 bp in the ancestors of Aves, Neognathae and Neoaves (~0.3, 0.4 and 0.2 Mb per My, [Fig S12 of 40]). Together, these observations suggest that genome contraction prior to the evolution of flight in the common ancestor of birds and bats must have occurred through relatively large chromosomal deletion events, but not through an increased rate of microdeletions.

Genome size variation between bird species has been linked to variation in metabolic cost of powered flight, with hummingbirds exhibiting the highest metabolism and smallest genomes [9, 92, 93], while flightless ratite birds display the largest genomes [2, 45, 94]. Our results lend further support to this connection between metabolic rate and constraint in genome size. We found that bird lineages that have lost flight (penguins and ostrich) are characterized by midsize deletions rates significantly lower than those of flying birds (2.3-fold on average; ks test, p = 0.0036; Figure 6, Dataset S3). This trend is also consistent with the results of a recent study indicating that TE removal through ectopic recombination occurs at a faster rate in the zebra finch (flying bird) than in the chicken (ground-dwelling bird) [42]. Furthermore, we observe that flightless bird lineages (penguins and ostrich) have gained generally less DNA during evolution than flying birds (Figures 3 and S1, Dataset S1). Thus, the larger genomes of flightless birds do not appear to reflect increased DNA gains, but slower removal of DNA relative to flying birds. In other words, the genomes of flightless birds are less dynamic overall than those of flying species.

In addition to their connection with powered flight, resting metabolic rates are correlated with body mass in mammals [95] and birds [96]. Interestingly, in our dataset we note that animals larger than other species within the same order (e.g. elephant vs. manatee and tenrec, cow vs. sheep, ostrich vs. tinamou) display lower micro- and midsize deletion rates (Figures 5 and 6). Similarly, megabats have larger body mass than microbats, and show a lower DNA loss coefficient (Figure 2). These observations are consistent with a relationship between body mass, resting metabolic rates, and genomic deletion rates. However, we did not detect any general correlation between body mass and DNA loss when all mammals and birds in our dataset are considered (Datasets S1 and S3), suggesting that there is no simple relationship between these parameters.

Finally, our results in bats are also consistent with the hypothesis that the metabolic requirements for powered flight constrain genome size [91, 97] (Figure 1). We found that bats have a DNA loss/gain ratio ~4.3-fold higher than the other mammals examined (Figure 2), as well as the highest midsize deletion rates (Figure 6, Dataset S3). Importantly, however, we observe that neither bats (Figures 5A and 6), nor flying birds (Figure 6 and Dataset S3) exhibit increased microdeletion rates relative to their flightless outgroups, again implying a predominant role of large deletion events in keeping the genomes of the flying species particularly streamlined. Further studies are warranted to better characterize the molecular mechanisms underlying these large chromosomal deletions and their biological significance in amniote evolution.

## METHODS

### Genomic and biological data

Version of assemblies and species names are listed in supplementary datasets. Genome sequences in fasta format were recovered from UCSC for mammals (http://hgdownload.soe.ucsc.edu/goldenPath) and from ftp://climb.genomics.cn/pub/10.5524/100001_101000/101000/assembly/[40] for birds. Body mass data are from [95, 98, 99] for mammals, and from [100] for birds.

### TE annotation

For all mammals besides bats, RepeatMasker outputs were downloaded from http://www.repeatmasker.org/genomicDatasets/RMGenomicDatasets.htmlLibraries, RepeatMasker open-4.0.5 [48] ran with the repeat library release 20140131 from repbase [101]. For bats, we obtained the TE annotation with RepeatMasker open-4.0.5 (using -e ncbi), using a custom library based on TE consensus sequences available in the literature [89, 90, 102–106]. For birds, we used available RepeatModeler outputs of 44 bird species [107], ran RepeatModeler locally for the duck genome, manually curated selected repeats and complemented these libraries with all available avian TE annotations [32, 101, 108]. All sequences were merged in one unique library using a custom Perl script (ReannTE_MergeFasta.pl, available at https://github.com/4ureliek/ReannTE). We obtained TE annotations by running RepeatMasker open-4.0.5 on all genomes of interest with the same library (using -e ncbi).

### Determination of ancient versus lineage specific TEs

For mammals, we classified TE families as lineage-specific or shared between placental mammals (Table S1). We compiled data from Repbase and annotations of the RepeatMasker libraries [48, 101], complemented by our own orthology assessment (combination of BLAT, observation of the conservation tracks on UCSC, and orthology assessment with the following script: https://github.com/4ureliek/TEorthology). In birds, the majority of TEs belong to the CR1 superfamily [32, 40]. CR1 have been active at least since the common ancestor of birds, with always several subfamilies at the same time [109]. This is because one CR1 lineage survived from the many lineages of LINE present in the common ancestor of birds and crocodilians [110]. CR1 consensus sequences tend to be similar between ancient and recent families (e.g., families CR1-E and CR1-J across most of avian evolution [109], which creates mis-annotations in the genome using RepeatMasker. Therefore, we relied on substitution rates to split TE-derived DNA into lineage-specific or shared. We developed a Perl script (https://github.com/4ureliek/Parsing-RepeatMasker-Outputs/blob/master/parseRM.pl) to parse the raw alignment outputs from RepeatMasker (.align files). This allowed us to use the corrected percentage of divergence of each copy to the consensus from these .align files (accounting for the extremely high rate of mutations at CpG sites). In case of overlaps (when a position could be aligned to more than one consensus sequence), the smallest percentage divergence was chosen for that position.

### DNA loss calculation

DNA loss coefficients were calculated as in [33]. We estimated lineage-specific DNA loss coefficients *k* with *E* = *A e*^*-kt*^, where *E* is the amount of extant ancestral sequence in the species considered, *A* the total ancestral assembly size, and *t* the time, leading to k = ln(A/X)/t). Assuming, for Eutherians, A = 2,800 Mb and t = 100 My, we get k = 0.0026 My^-1^ for human (X = assembly size minus gains = 2,150 Mb). See Dataset S1 for all values and coefficients of other species. DNA loss rates in Mb/My were obtained by dividing the amount of loss by the divergence time.

Note that changing the ancestral assembly size would affect the numbers but not the differences between species or the correlations. For example, using 2,600 Mb as in the mouse genome paper [30] would simply reduce loss of 200 Mb for each mammal. For birds, there is not a reconstruction of the ancestral assembly size of Neoaves as there is for mammals [111]. The ancestral genome size of Neoaves is estimated to range from 1.5 to 1.7 Gb. Therefore, we considered low and high boundaries of ancestral assembly sizes being 1.2 and 1.4 Gb, respectively, and we used the middle point of 1.3 Gb for all calculations. We also characterized total loss at the same evolutionary timescales as the ones of our microdeletions and midsize deletion calculations (Figure S4).

Additionally, with this method, we only measure net totals of DNA gain and loss amounts. Therefore, they likely represent underestimations of genome dynamics, even more so for the species with higher TE removal. For example, two species may have similar measured gains and loss, but one would in fact have higher efficiency of TE removal. The latter would appear as lower gains, which consequently translates in lower loss with our method. Such bias can be addressed by measuring DNA gain and loss at shorter evolutionary time scales and verify the extent of the gain and loss dynamics (Figure S4). Importantly, the results at shorter time scales recapitulate the ones of the 70 My scale, with the woodpecker and flightless bird (ostrich and penguins) lineages showing the most and the least dynamic genomes, respectively.

### Analysis of microdeletions using a multi species alignment

We developed custom Perl scripts (MAFmicrodel), available at https://github.com/4ureliek/MAF_parsing). In a first step, we extracted alignment lines corresponding to species of interest (mammals or birds) from the MultiZ alignment of human chromosomes 1 to 22 with 100 other species from the UCSC genome browser (MAF format, http://hgdownload.cse.ucsc.edu/goldenPath/hg19/multiz100way/); see the usage of the MAF_microdel--1--get-gaps.pl script. Studied species are listed on Figure 5, with *Monodelphis domestica* and *Anolis carolinensis* as outgroups for mammals and birds, respectively (Dataset S2). Gaps were then extracted and listed in bed format for blocks containing alignment information for all species. For this analysis, we also filtered out blocks shorter than 50 bp (MAF_microdel--1--get-gaps.pl). In a second step (scripts MAF_microdel--2--analyze-gaps-XXX.pl), gaps in alignments were placed on a phylogenetic tree based on parsimony and using intersections of gap coordinates with Bedtools [112]. To limit biases arising from counting specific gaps occurring at the same location as shared, a gap is considered shared between two species only if the reciprocal overlap is >80% (options -f 0.80 and -r of intersectBed). The deletion rate is then calculated by dividing the amount of gaps strictly specific to each branch (not present in any of the other species) by the length of each branch (in My).

### Analysis of microdeletions and midsize deletions for trio of species

We developed custom Perl scripts for the analysis of microdeletion and midsize deletions in species trios, available at https://github.com/4ureliek/DelGet. Most parameters can be adjusted through the configuration file. The steps are as follow: (i) Pick random positions in the outgroup species (ii) Extract X bp of sequences from this species’ randomization, separated by Y bp. We chose X = 100 bp with Y = 10 kb (because the N50 of the contigs in some assemblies are <20 kb). (iii) Use BLAT to obtain hits in other species. Hits are kept only if (a) they are both on the same scaffold of the target; (b) both hit lengths are >120 bp; (c) both hits are on the same strand, score of best hit/next hit is >0.9 (to avoid uncertainty related to repeats), and (d) the region does not overlap with gaps in assembly (‘N’ nucleotides). (iv) The sequences between anchors are extracted for all three genomes. (v) These sequences are aligned with MUSCLE [113]. (vi) Gaps in alignments are placed on a phylogenetic tree through intersections with Bedtools [112]. A gap is considered shared between 2 species if the reciprocal overlap is >85% (option -f 0.85 and option –r for reciprocity). Additionally, when only one base interrupts a gaps (e.g. GTGC---------A------ATGTC) it is skipped and the 2 gaps are merged in one (its length being corrected by 1 bp).

### Screening for large deletions

Using a custom Perl script (maf_get_large_indels.pl, available at https://github.com/4ureliek/MAF_parsing), we recovered coordinates of indels >1 kb for each species in the MultiZ alignment of human chromosomes 1 to 22 with 100 other species from the UCSC genome browser (MAF format, http://hgdownload.cse.ucsc.edu/goldenPath/hg19/multiz100way/). The script outputs two types of large indels: first, a list of indels corresponding to cases where the sequence before and after is contiguous (“C” lines), implying that this region was either deleted in the source or inserted in the reference sequence (or a combination of both). This will lead to underestimating event lengths in case of shared repeats, but to avoid overestimating indel lengths due to TEs inserted in the reference (human), repeated sequences are discarded (as annotated by RepeatMasker and soft-masked in lower cases in the alignment). The script also outputs a list of indels corresponding to all consecutive empty data for a given species (no data on the browser, or double line when there are non aligning bases; these gaps could be shared between several species) only when the empty data is interrupting a scaffold in the species of interest. When there are inserted sequences in the species of interest, the script only outputs cases where the length in the reference (no lowercases) minus the insertion length is higher than the minimum length specified (here, 1 kb). Note that these gaps could arise from mis-assemblies or from segmental duplications in the reference, and all of them would require to be validated prior to any interpretations.

## AKNOWLEDGMENTS

We thank Aditi Rambani for her contribution in designing the ‘DelGet’ pipeline to find midsize deletions. We are grateful to Xiaoyu Zhuo, Zev Kronenberg, Carson Holt, Barry Moore, and Mark Yandell for their help with bioinformatics and statistics, and to Rachel Cosby for helpful discussions. We further thank Cai Li for providing avian neutral substitution rates, Benoit Nabholz and Claudia C. Weber for providing avian body mass data, and Lel Eory and David Burt for providing RepeatModeler libraries. This work was supported by grant R01GM077582 to C.F. from the National Institutes of Health.

## SUPPLEMENTARY MATERIAL

Dataset S1: gain and loss of DNA (data of Figures 2, 3, 4 S1 and S4)

Dataset S2: Analysis of microdeletions using a multi species alignment (data of Figure 5)

Dataset S3: Analysis of microdeletions and midsize deletions for trio of species (data of Figure 6)

Dataset S4: Statistical significance for the pairwise comparisons of distributions of Figures S2, S3 and S5

**Figure S1:**
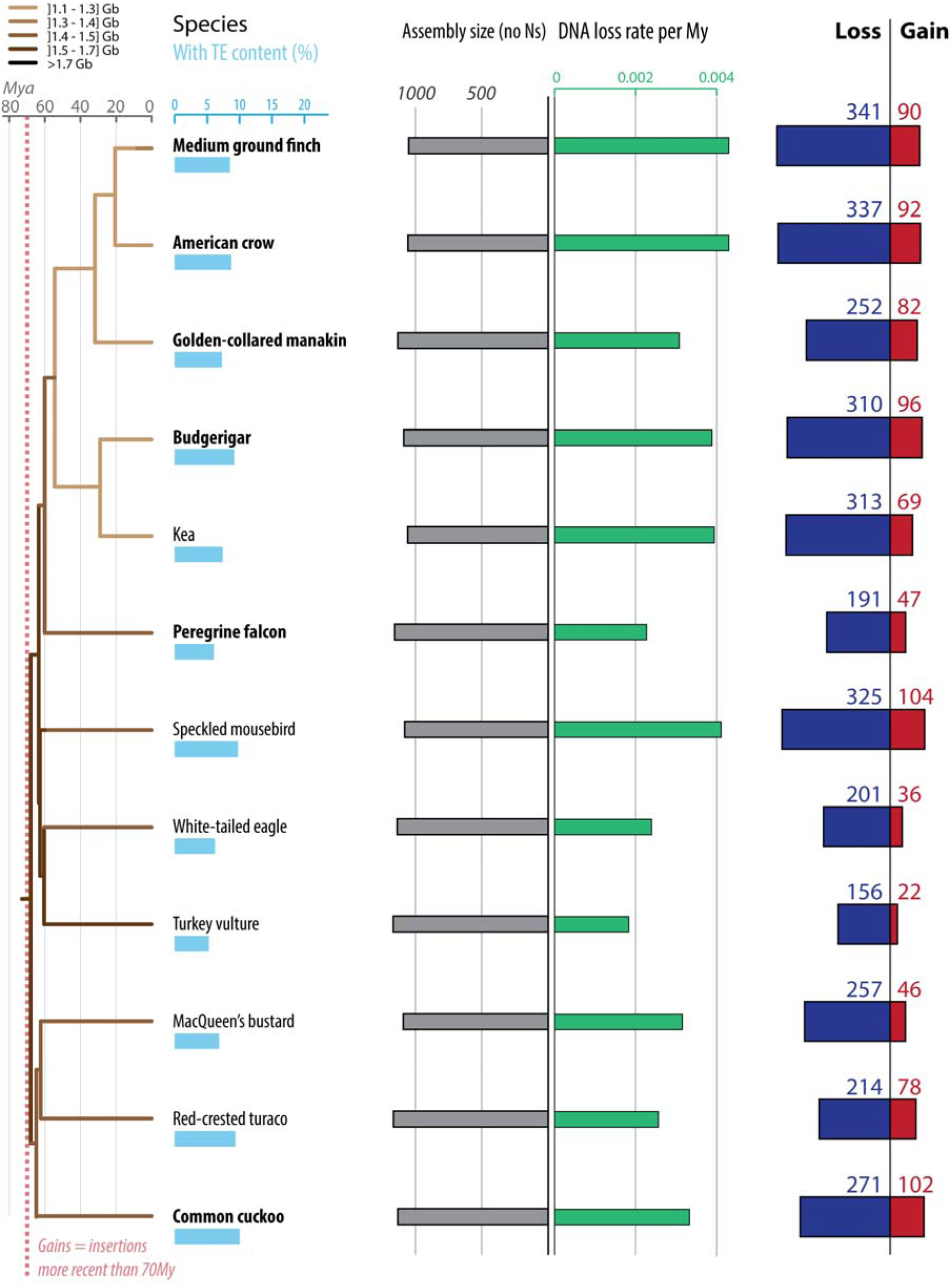
Gain and loss of DNA in 12 additional birds. Figure reads exactly as Figure 3, with different species represented. For each species, phylogenetic relationship (left panel) [49], TE content (light blue bars), assembly sizes (with N removed, grey bars), DNA loss rates per My (green bars) as well as gain (red) and loss (dark blue) of DNA are shown. Species names in bold correspond to high-coverage genomes, and the others to low-coverage genomes [40]. DNA gain corresponds to TE insertions younger than 70 My. DNA loss amounts and rates are calculated as in [33] using a common ancestor size of 1,300 Mb (Methods). Phylogenetic tree is color coded based on genome sizes in pg (combination of data from [114] and extrapolations from assembly sizes and coverage), based on parsimony as in [52]. See Dataset S1 for numbers, calculation steps and assemblies details. All numbers are in Mb.

**Figure S2:**
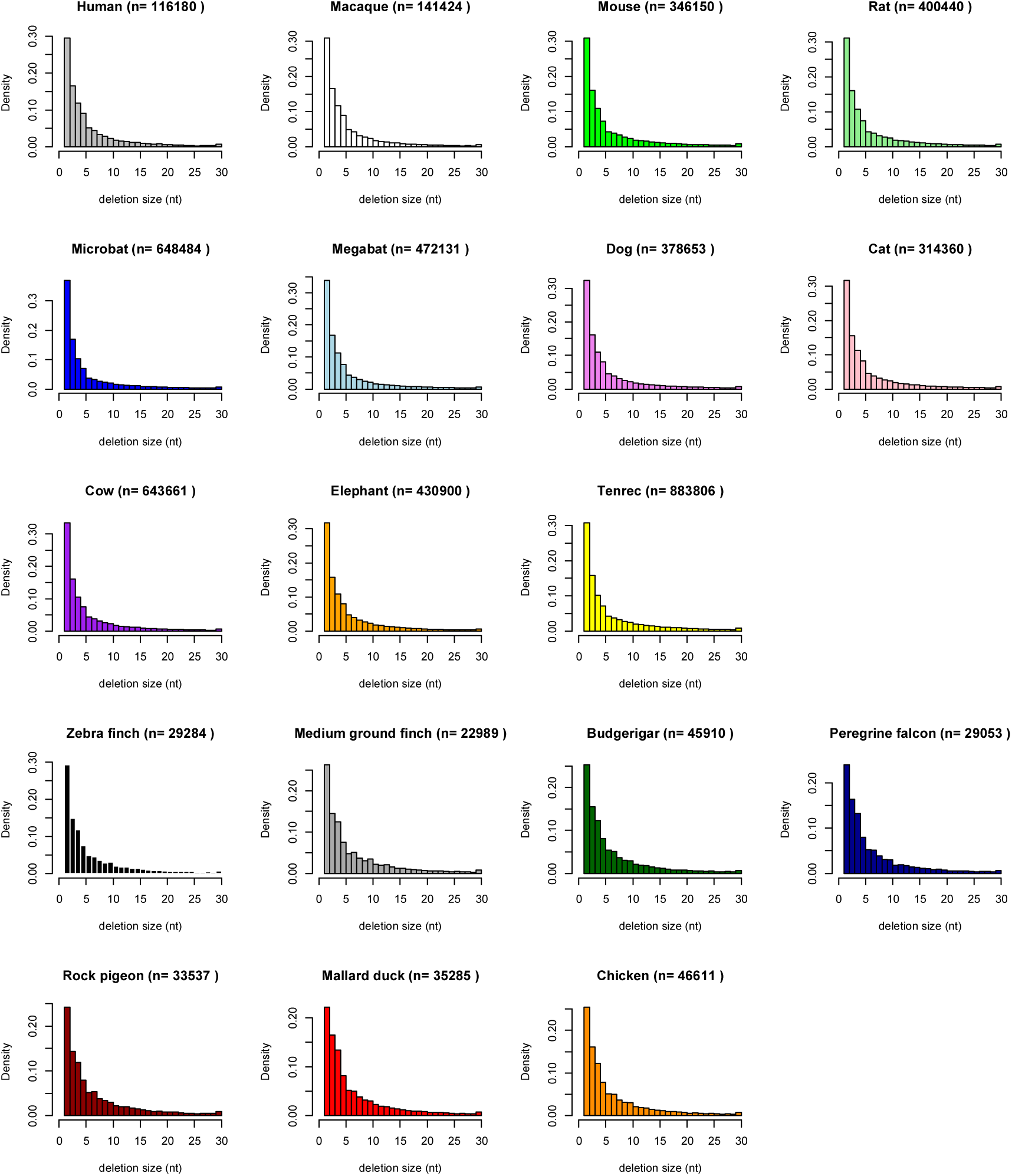
Microdeletion spectrums of mammals and birds. The data plotted is that of Figure 5, for the terminal branches (gaps specific to each species). The histograms are done in R [117], with for example for human: hist(hs$V1, freq=F, right=F, border="black", col="grey", main=paste("Human (n=",length(hs$V1),")"), xlab = "deletion size (nt)", xlim=c(0,30),breaks=30), xlim=c(0,30), breaks=30)), where hs corresponds to the list of gap lengths specific to human. Statistical significance is based on 1,000 replicates of Kolmogorov–Smirnov tests on samples of 5,000 values: two distributions are considered significantly different when at least 950 out of 1,000 tests have *p* < 0.05. Data and R command lines can be found in Dataset S4, sheet ‘micro_del’. Densities and not frequencies are plotted to allow comparison between species. The number of deletion events (inferred from gaps in the alignment) is listed on top of the histograms (n).

**Figure S3:**
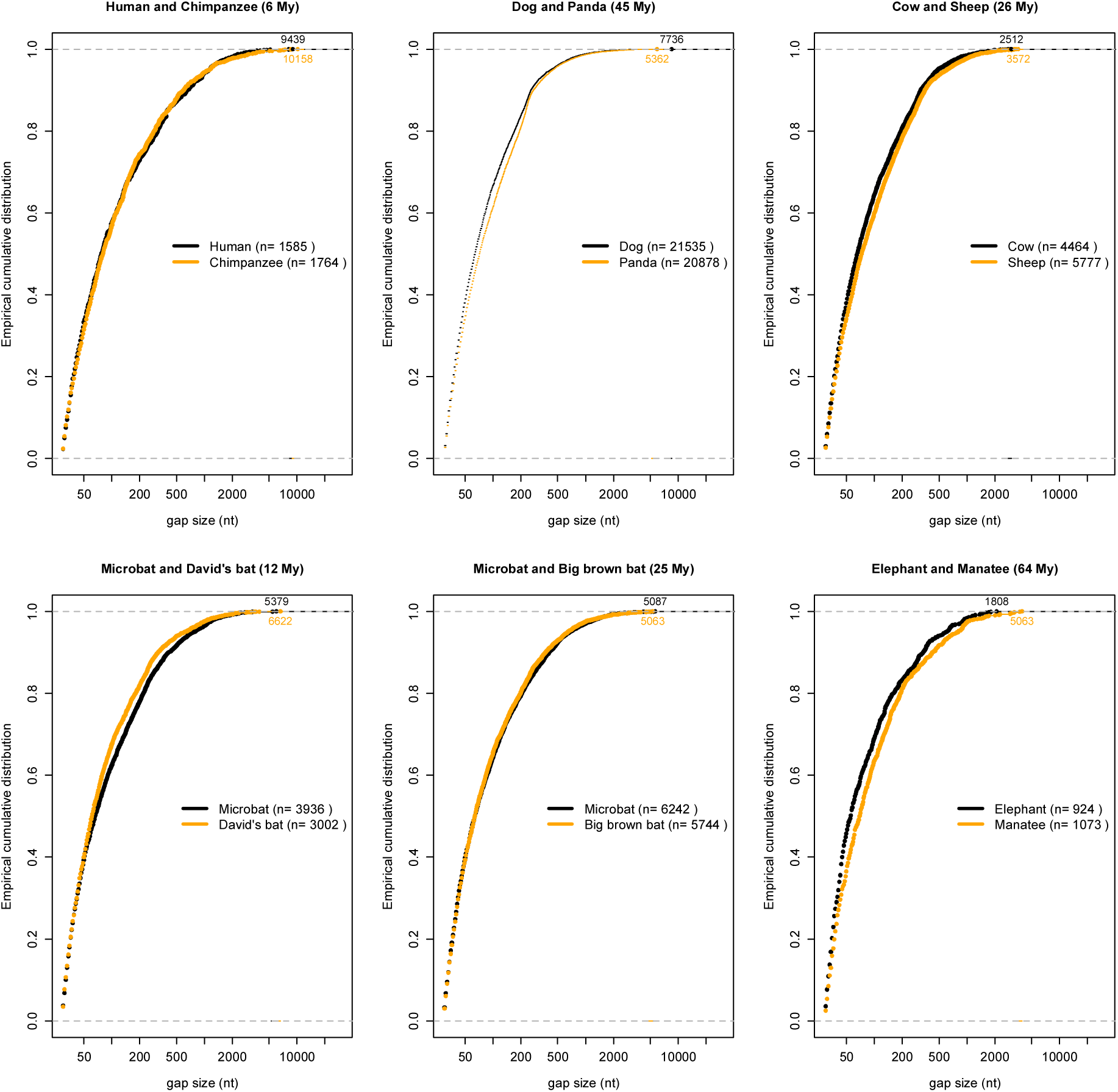
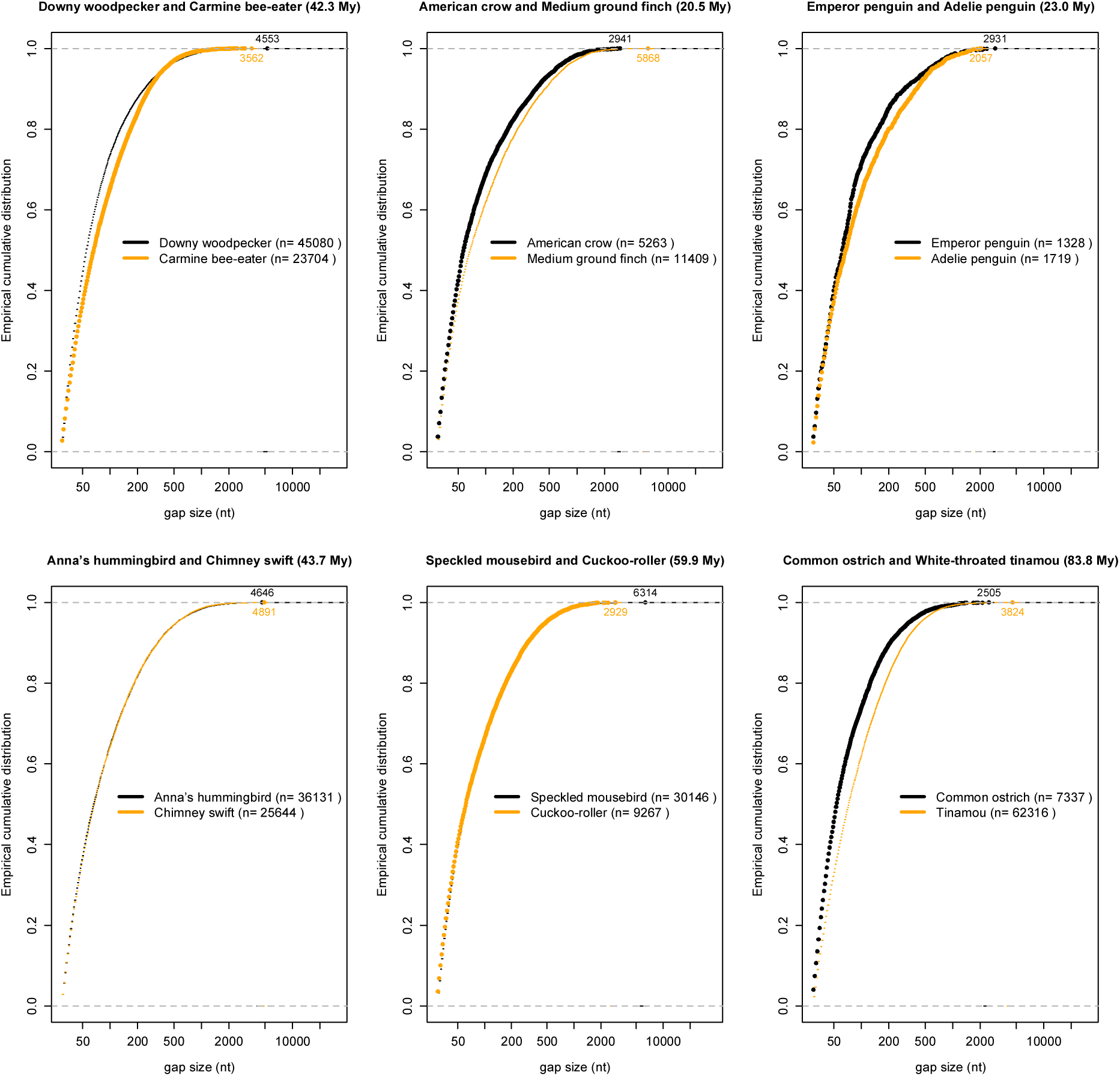
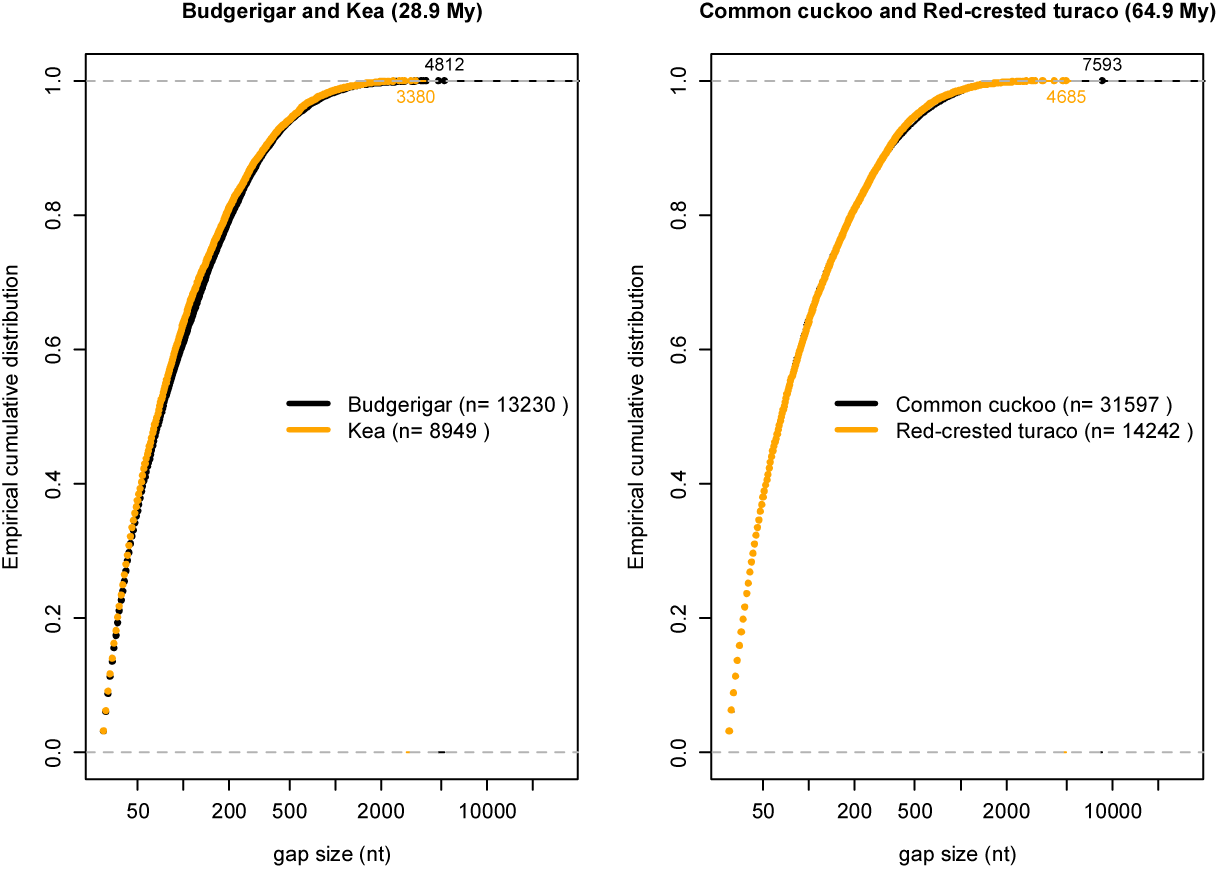
Midsize deletion spectrums of mammals and birds. A. Mammals (data of Figure 6). B. Birds (data of Figure 6). C. Four additional birds. The empirical cumulative distributions (ecd) are done in R [117] (command lines can be found in Dataset S3), and statistical tests are done as in Figure S2 (detailed in Dataset S4). Note the log scale for the x axis, for better visualization. On ecd graphs, differences in the slopes of the curves reveal differences in density: for example, the elephant has more deletions of ~200 bp than the manatee. Indeed, such representation (cumulative) allows to directly visualize the proportion of events under a certain size; for example, between 10 and 23% of the deletion events recovered in each species were larger than 200 bp. Clearly distinct curves suggest significant differences, but the dog and the panda distributions are the only ones that have enough events to be found significantly different (Dataset S4). For C (four additional birds), the DelGet pipeline (Methods) was run for the budgerigar and kea with the American crow as outgroup (155.1 Mb of alignment), and for the common cuckoo and the red-crested turaco with the Chuck-Will’s-widow as outgroup (128.5 Mb of alignment). Resulting microdeletion and midsize deletion rates are intermediate to those of ostrich and Anna’s hummingbird (Dataset S3). The largest events are labeled on the graphs and have been validated *in silico* through careful manual examination of the alignments. Additionally, their length has been corrected in case of lineage-specific TE insertions in the other species. Breakpoints are as follow for mammals: human (hg38 chr10:81816824), chimpanzee (panTro4 chr14:92050068), dog (canFam3 chr20:33042110), panda (ailMel1 GL194157.1:120088), microbat (myoLuc2 GL429820:6762144 for group I and GL429772:17082212 for group II), Brandt’s bat (ASM41265v1 gb|KE161863.1|:221999), big brown bat (Eptesicus_fuscus_assembly1.0 gb|ALEH01017523.1|:41408), cow (bosTau8 chr22:10255818), sheep (oviAri3 chr11:31801480), manatee (TriManLat1.0 gi|460713488|ref|NW_004444110.1|:1177735), elephant (loxAfr3 scaffold_6:35237920). For birds, assemblies are as in [40] (Dataset S1) and breakpoints are as follow: downy woodpecker (scaffold783:2013016), carmine bee-eater (scaffold45549:18816), American crow (scaffold116:6801125), medium ground finch (scaffold40:8463065), emperor penguin (Scaffold500:719123), Adelie penguin (Scaffold241:3188990), Anna’s hummingbird (scaffold104:965959), chimney swift (scaffold114:243360), speckled mousebird (scaffold41959:22764), cuckoo-roller (scaffold18151:34945), common ostrich (scaffold638:67481), tinamou (scaffold12594:11743), budgerigar (scf900160277013:1211857), kea (scaffold13226:14568), common cuckoo (scaffold492:288210), red-crested turaco (scaffold3767:111605).

**Figure S4:**
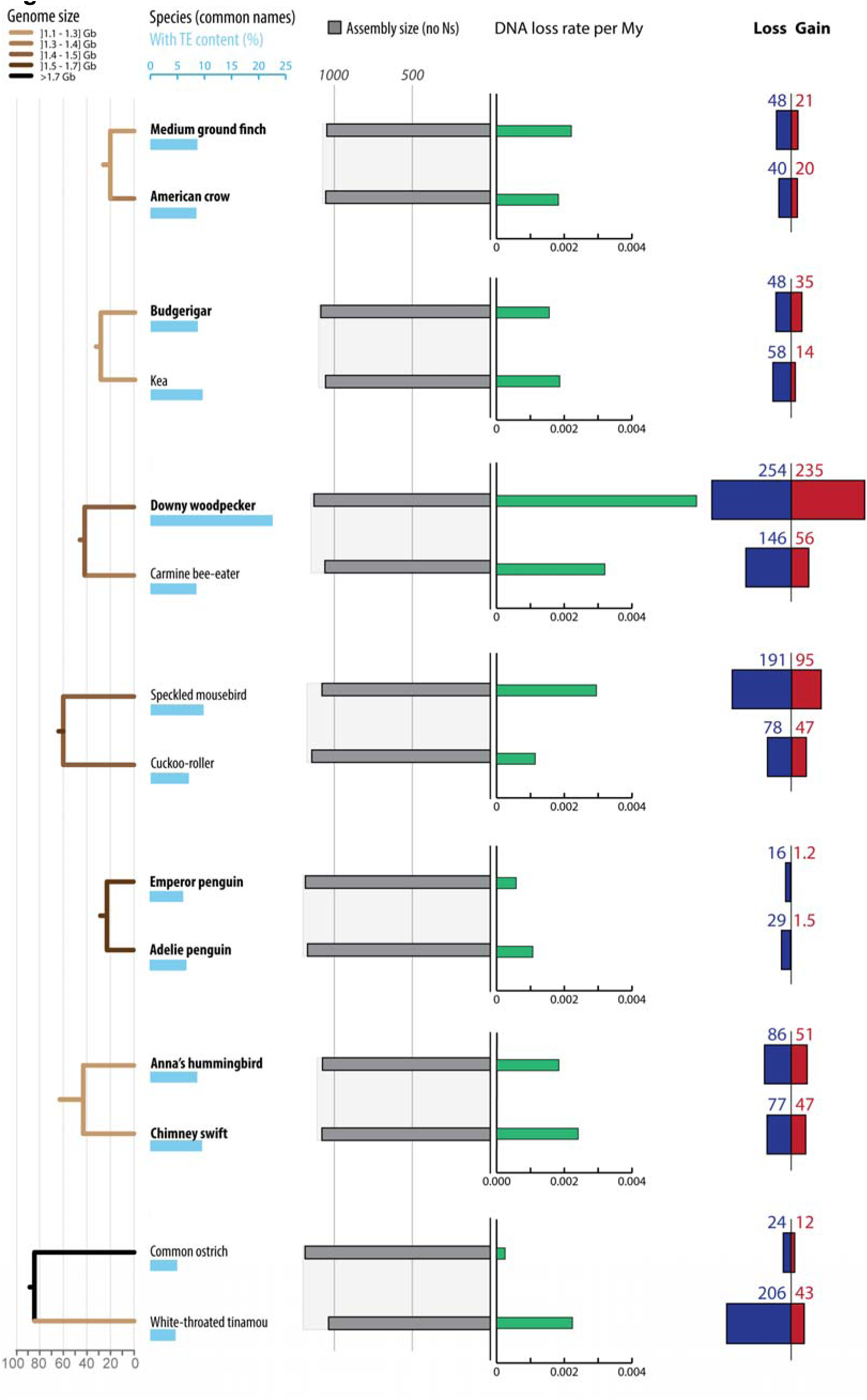
Gain and loss of DNA for seven duos of birds. Different species or different time scales than Figures 2 or S1 are considered, to match the times of microdeletion and midsize deletion rates of Figure 6. For each duo, we calculated the amounts of gain and loss of DNA after the two birds diverged. DNA gains (in red) correspond to lineage-specific TEs since the split of the last common ancestor of the three birds considered. Ancestral assembly size used for calculation of DNA loss (Methods) for each duo is shown with light grey highlight behind assembly sizes. It is based on the common ancestor genome size, which was estimated using the genome sizes of all species (combination of data from [114] and extrapolations from assembly sizes and coverage, color coded on the branches) and based on parsimony [52]. Estimations of the ancestral assembly size could be flawed and affect DNA loss estimations; but we observe that microdeletion and midsize deletion rates still mildly correlate with the DNA loss coefficients when calculated at the same time scale (R^2^ = 0.52 with *p* = 0.027 and R^2^ = 0.44 with *p* = 0.087 respectively; R^2^ = 0.51 with *p* = 0.046 for both rates combined). Importantly, DNA gain and loss are still significantly correlated (R^2^ = 0.82 with *p* = 0.0003, Dataset S1), which is indicating of a continuous counteraction of DNA gain by DNA loss. For each species, phylogenetic relationship (left panel) [49], TE content (light blue bars), assembly sizes (with N removed, grey bars), DNA loss rates per My (green bars) as well as gain (red) and loss (dark blue) of DNA are shown. Species names in bold correspond to high-coverage genomes, and the others to low-coverage genomes [40]. See Dataset S1 for numbers, calculation steps and details. All numbers are in Mb.

**Figure S5:**
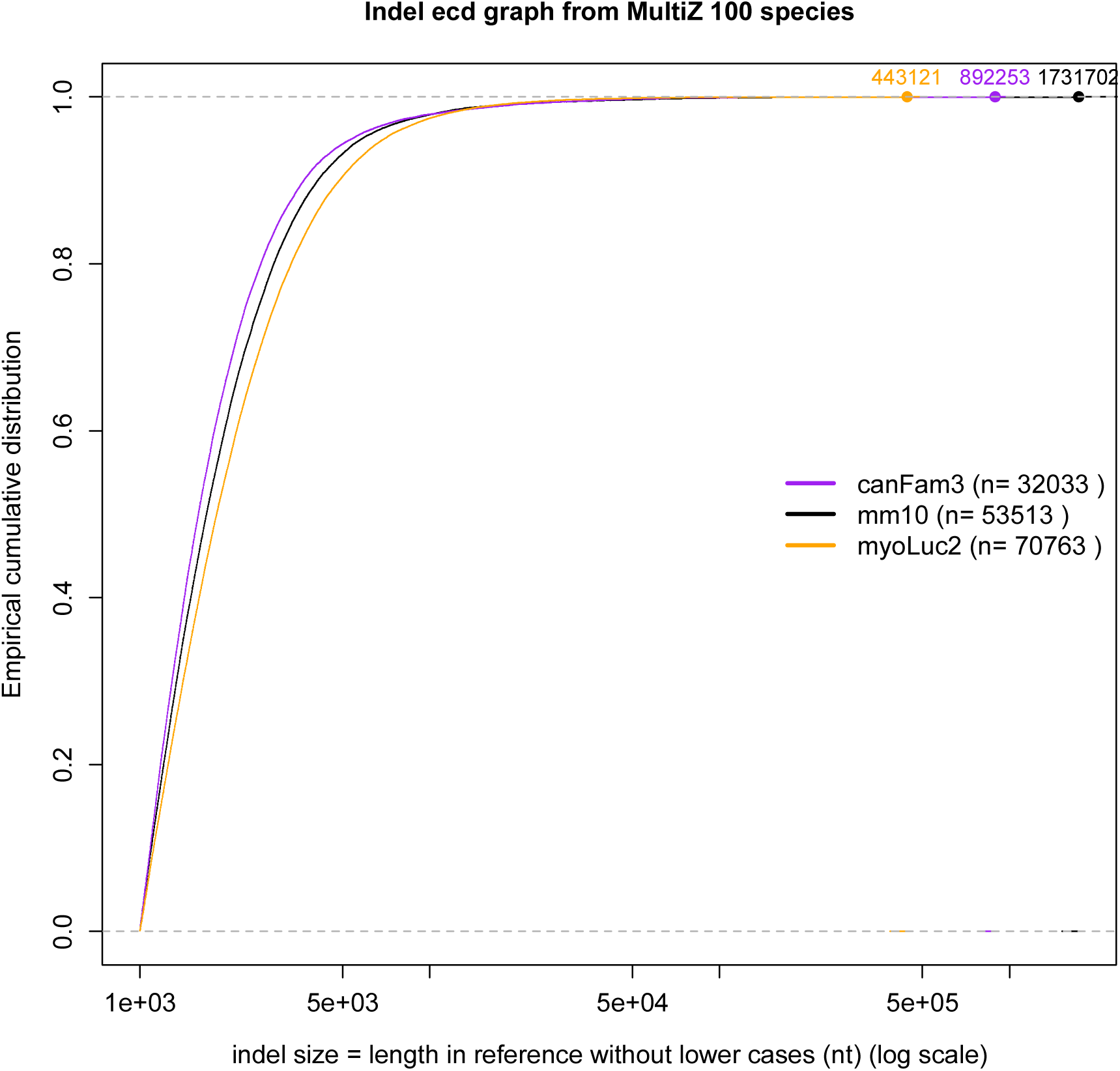
Size distribution of large indels in mammals. Large indels screened from the MultiZ 100-species alignment from UCSC genome browser (“empty data”, see Methods). The empirical cumulative distributions are done in R [117], as in Figure S3. Statistical tests are done as in Figure S3 and are detailed in Dataset S4. Largest events are labeled on the graphs, and we verified that the three corresponding sequences in hg19 could not be found in the query species (blastn against whole-genome shotgun contigs, with repeats masked). For example, the largest deletion of the microbat is 443,121 bp long in hg19 (976,711 bp with lowercases) and is at position GL430240:588646 in myoLuc2, corresponding to chr1:238471164-239447875 in hg19. This event seems shared with other *Myotis* species and with the big brown bat *Eptesicus fuscus*, but not with the two megabats.

**Table S1:**
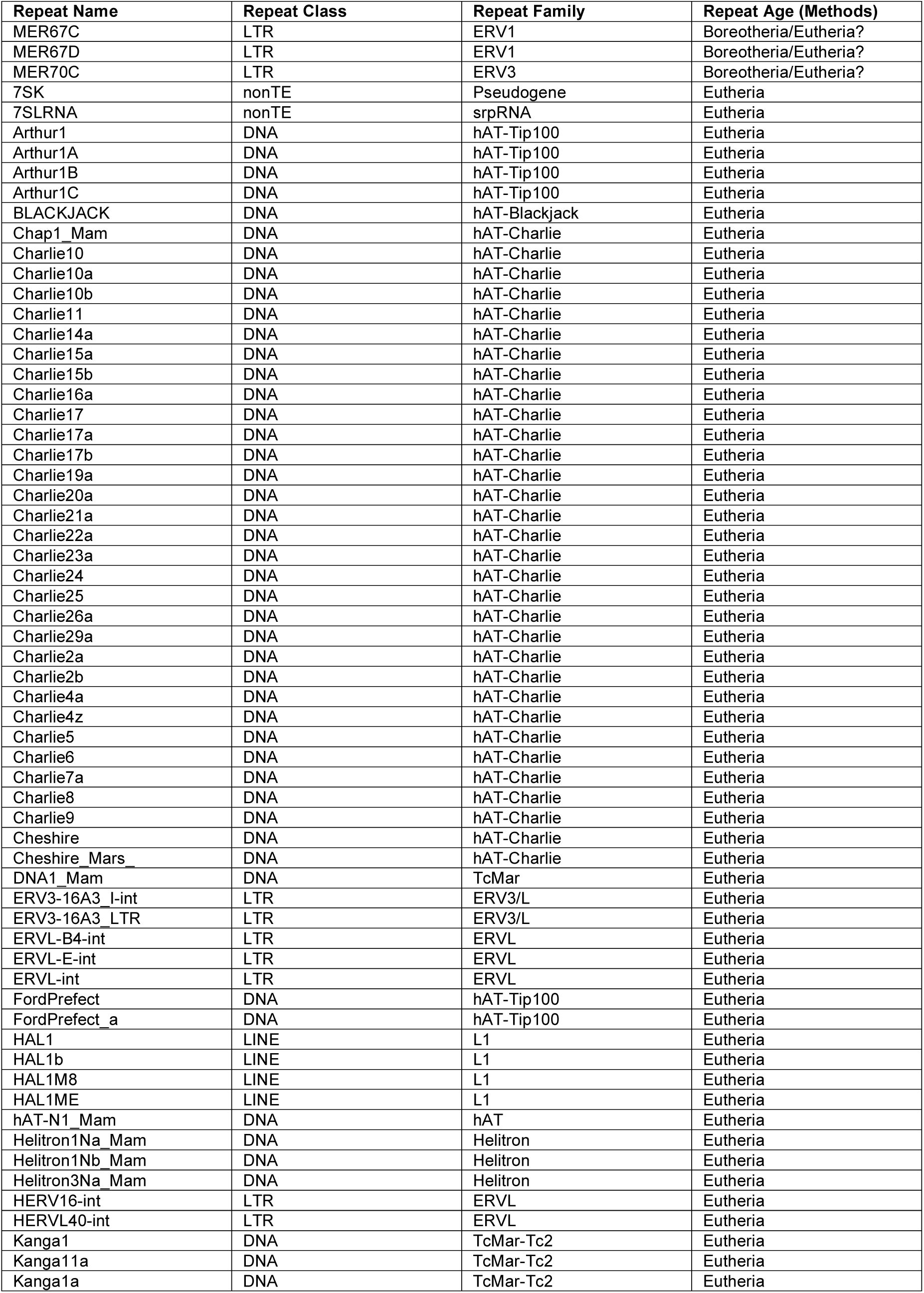
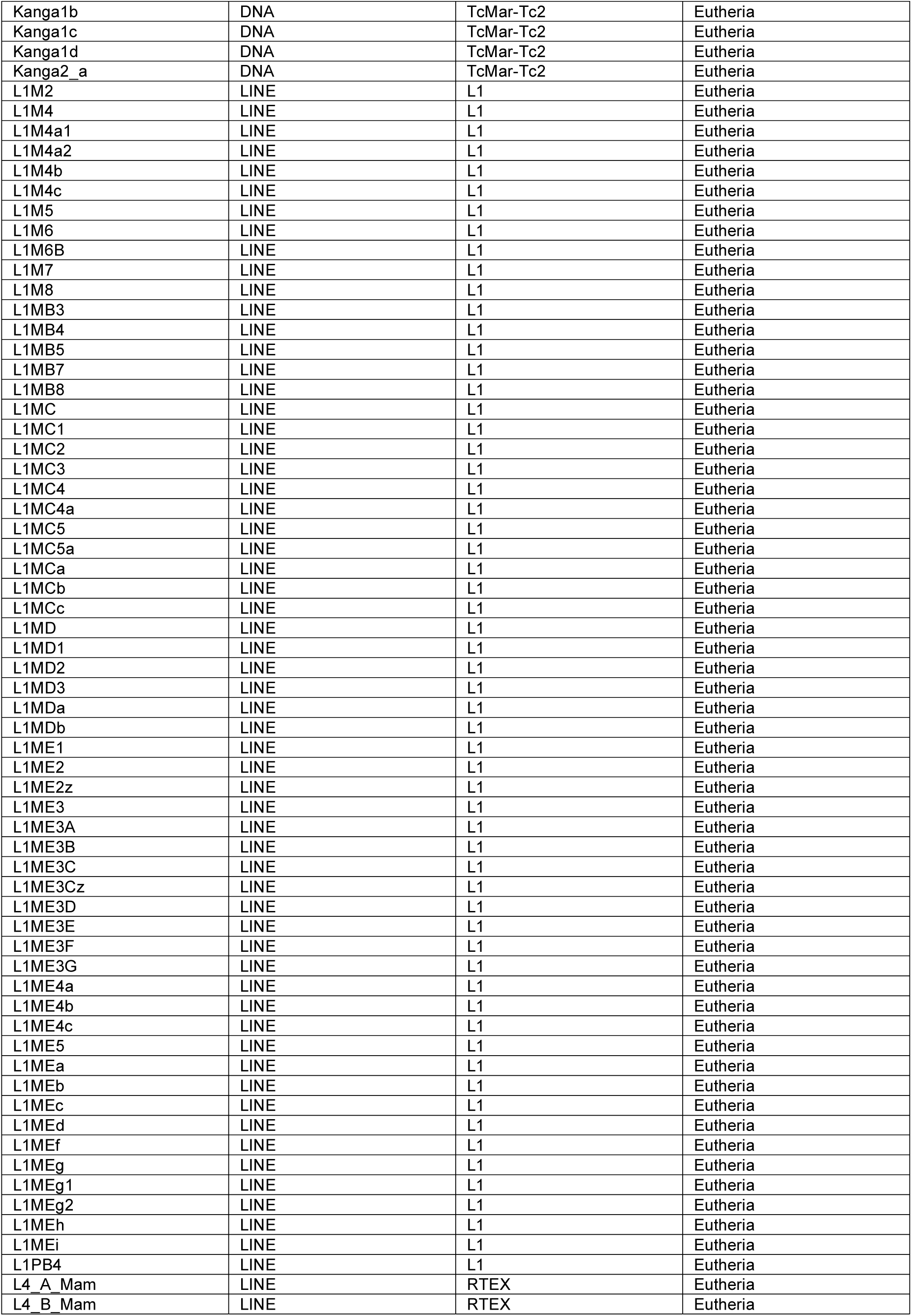
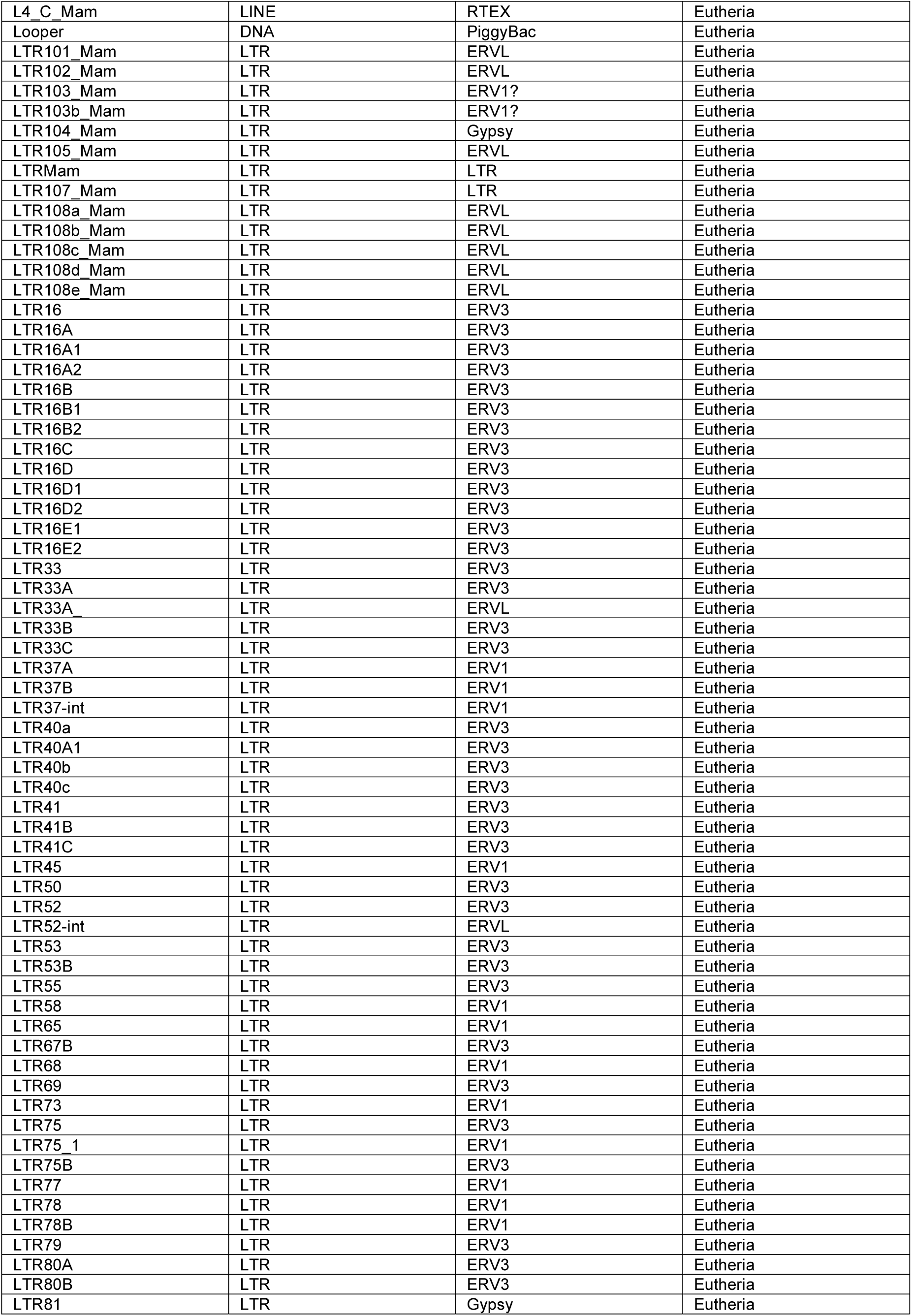
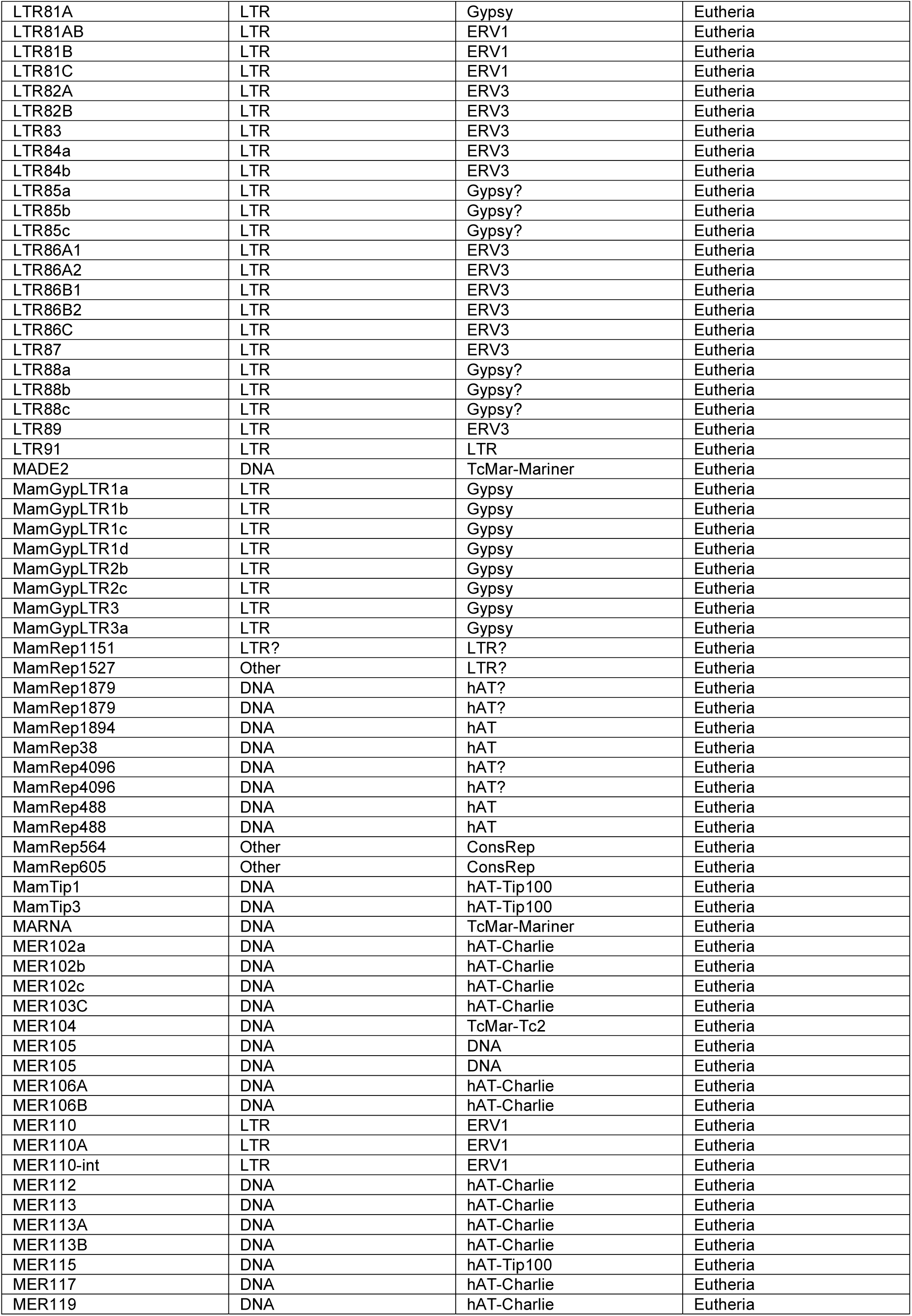
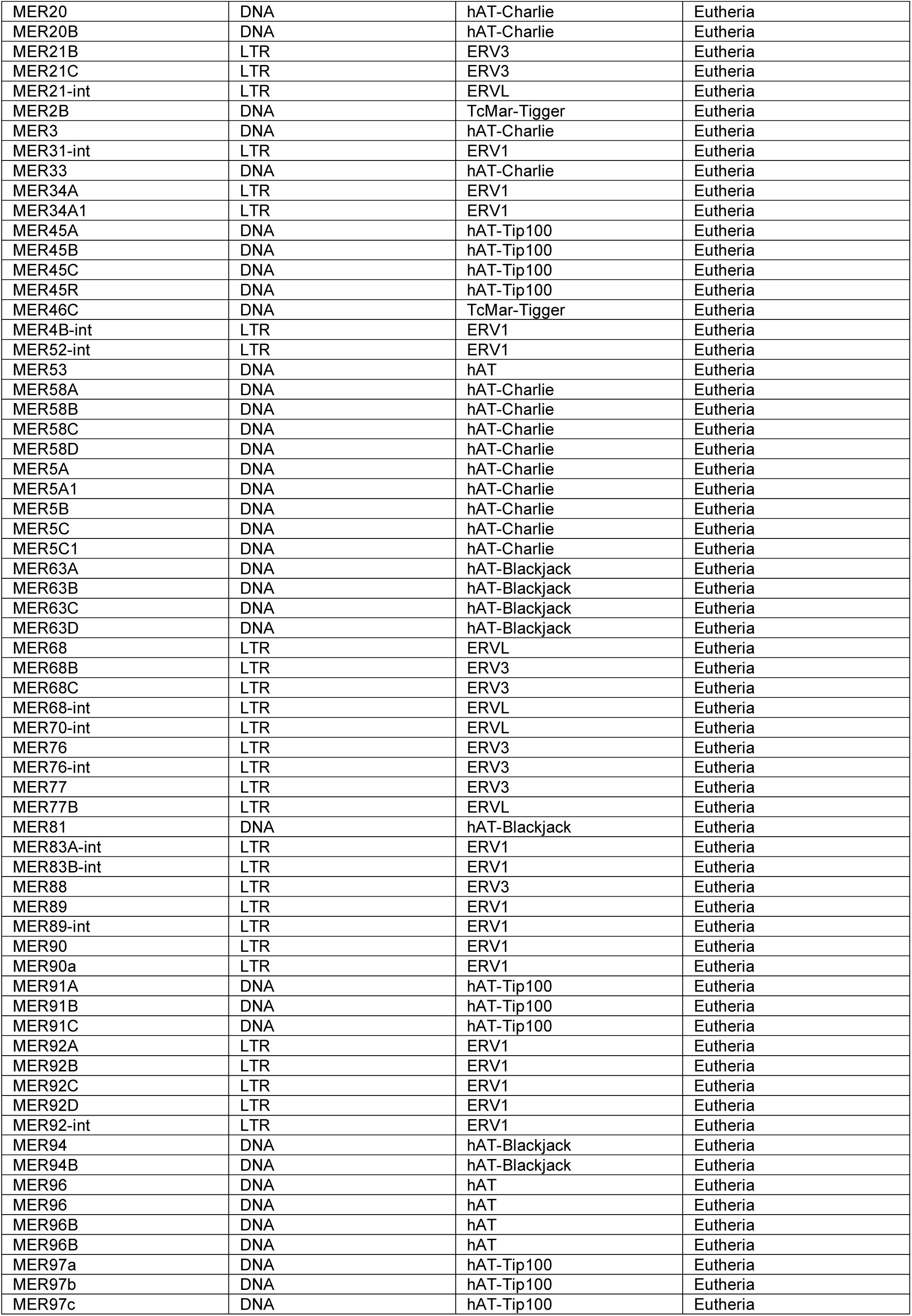
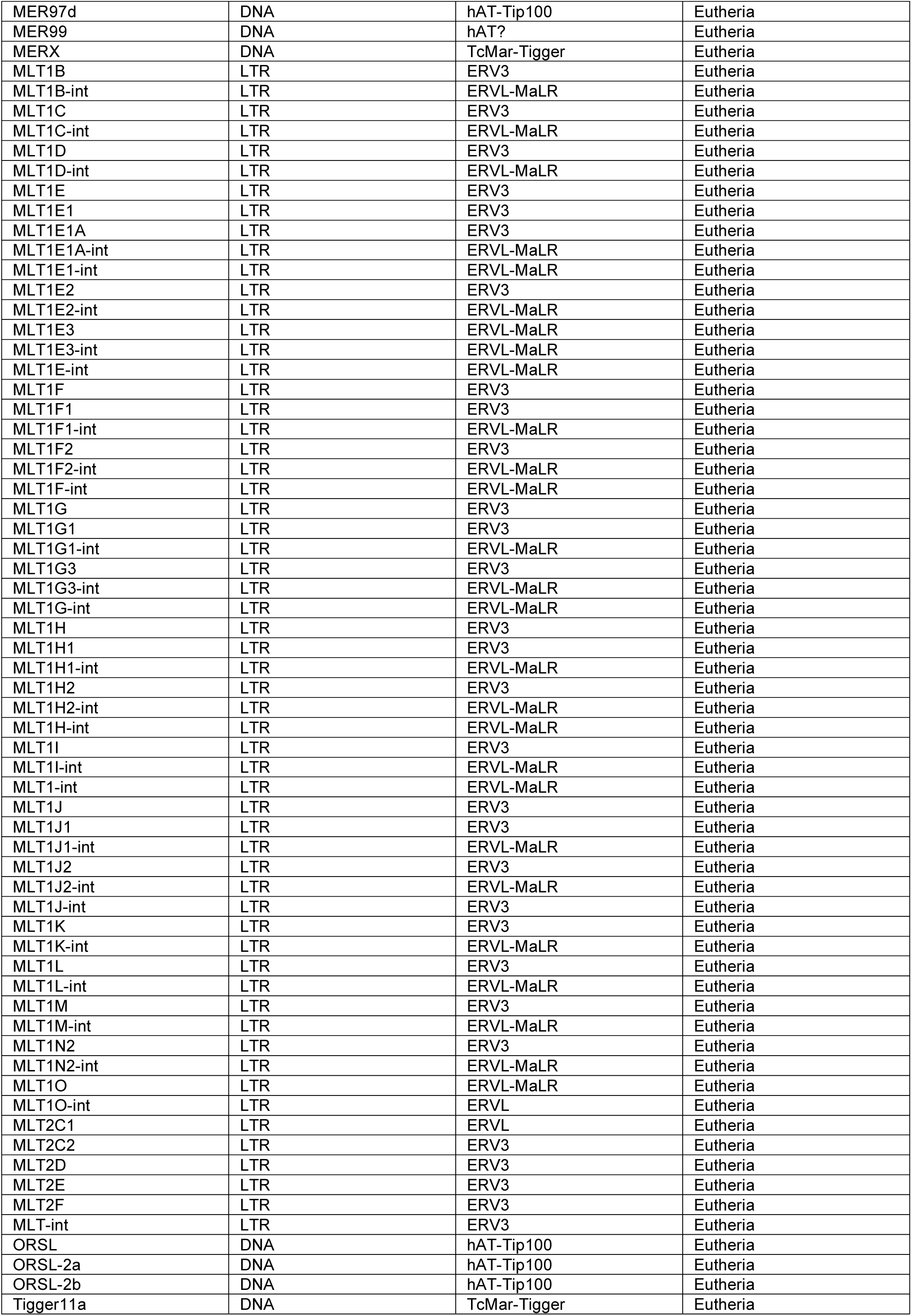
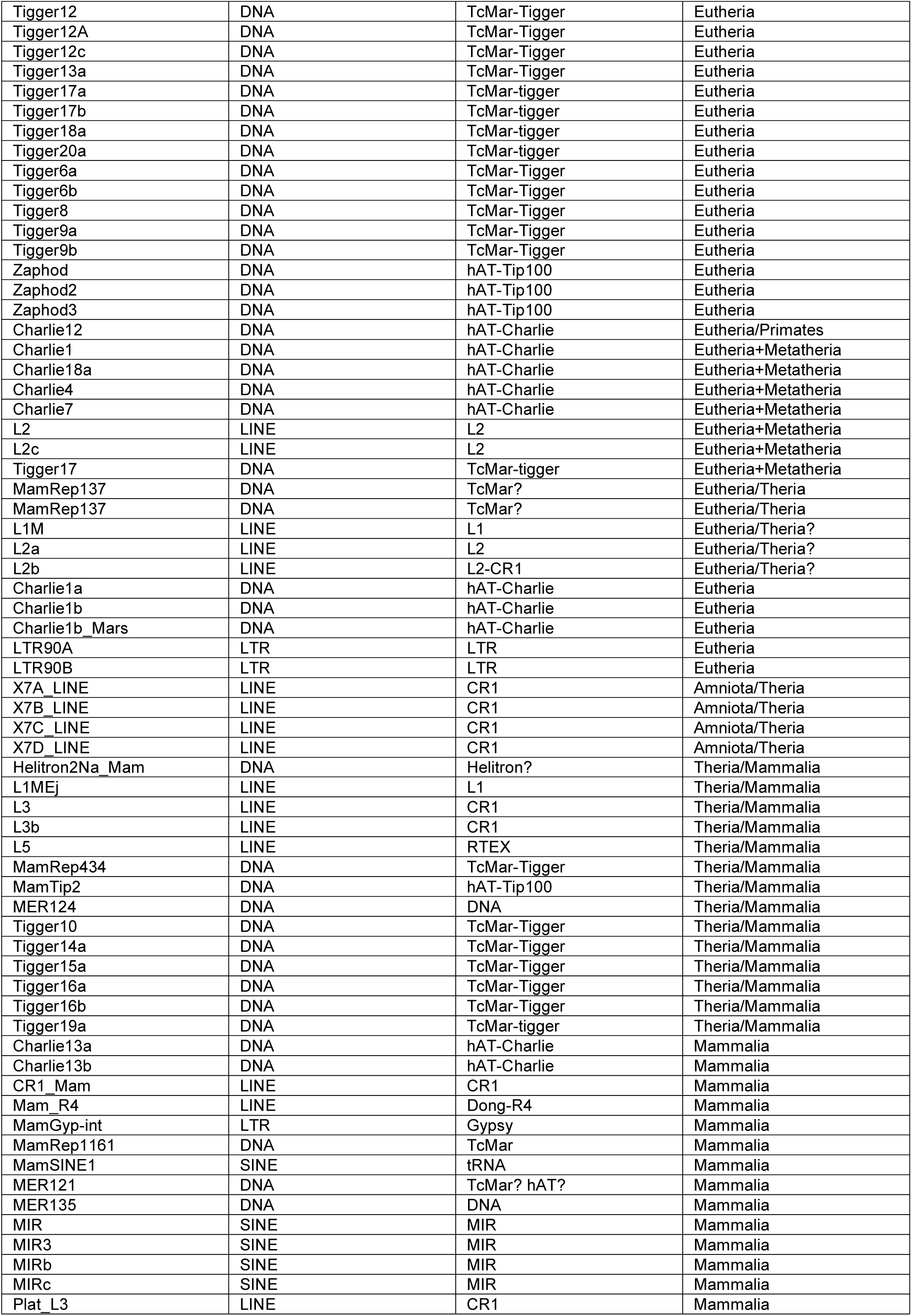
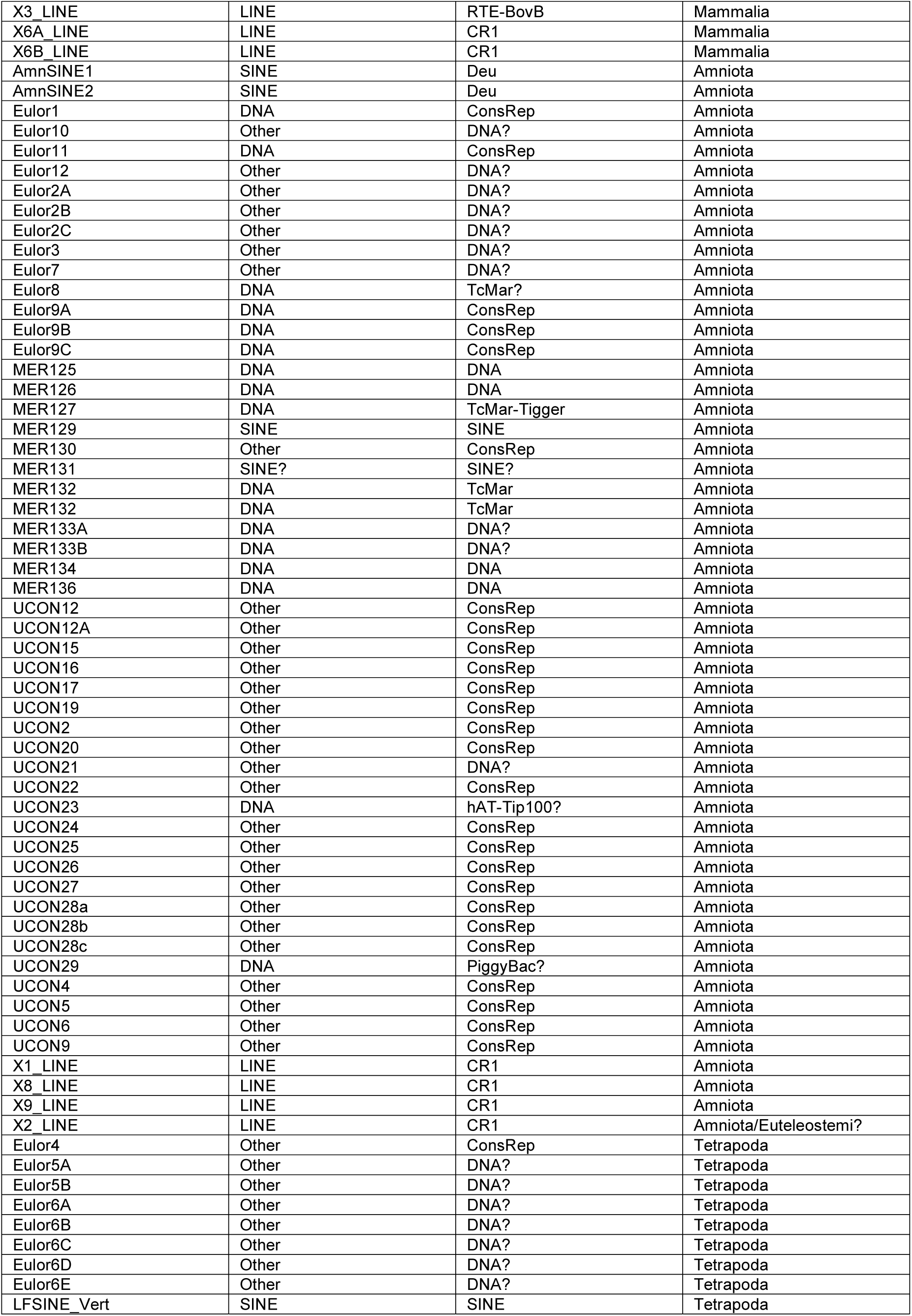
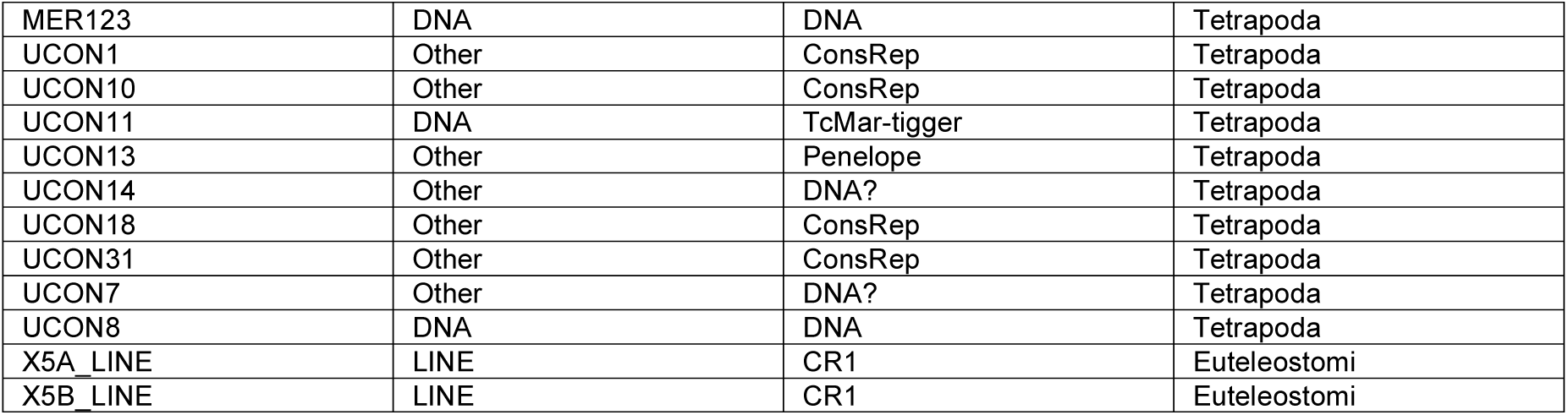
List of ancient mammalian TEs

